# A nucleus-forming jumbophage evades CRISPR-Cas DNA targeting but is vulnerable to type III RNA-based immunity

**DOI:** 10.1101/782524

**Authors:** Lucia M. Malone, Suzanne L. Warring, Simon A. Jackson, Carolin Warnecke, Paul P. Gardner, Laura F. Gumy, Peter C. Fineran

## Abstract

CRISPR-Cas systems provide bacteria with adaptive immunity against bacteriophages^1^. However, DNA modification^2,3^, the production of anti-CRISPR proteins^4,5^ and potentially other strategies enable phages to evade CRISPR-Cas. Here we discovered a *Serratia* jumbophage that evaded type I CRISPR-Cas systems, but was sensitive to type III immunity. Jumbophage infection resulted in a nucleus-like structure enclosed by a proteinaceous phage shell – a phenomenon only reported recently for distantly related *Pseudomonas* phages^6,7^. All three native CRISPR-Cas complexes in *Serratia* – type I-E, I-F and III-A – were spatially excluded from the phage nucleus and phage DNA was not targeted. However, the type III-A system still arrested jumbophage infection by targeting phage RNA in the cytoplasm in a process requiring Cas7, Cas10 and an accessory nuclease. Type III, but not type I, systems frequently targeted nucleus-forming jumbophages that were identified in global viral sequence datasets. These findings explain why many bacteria harbour both RNA- and DNA-targeting CRISPR-Cas systems^1,8^. Together, our results indicate that jumbophage nucleus-like compartments serve as a barrier to DNA-targeting, but not RNA-targeting defences, and that this phenomenon is widespread amongst jumbophages.

---

CRISPR-Cas systems consist of array(s) with invader-derived spacers separated by short repeats and the Cas (CRISPR-associated) proteins that provide the enzymatic machinery for immunity^1^. During phage infection, invading genetic material can be acquired into CRISPR arrays as new spacers^9,10^. Expression and processing of the CRISPR array(s) results in crRNA guides, onto which Cas proteins assemble, forming surveillance complexes^11,12^. In the interference step, recognition of foreign genetic material complementary to the crRNA leads to degradation of the phage nucleic acids and infection is arrested^13^. CRISPR-Cas systems are classified into two classes and six different types and bacteria often harbour multiple systems^8^. For example, *Serratia* sp. ATCC 39006 encodes three systems – type I-E and I-F that target DNA and type III-A that targets DNA and RNA^14^.

To identify new CRISPR-Cas evasion mechanisms, we isolated phages infecting *Serratia* and assessed their sensitivity to CRISPR-Cas immunity. Of these phages, a member of the *Myoviridae* family was selected for further characterisation and named PCH45 (Figure 1A). PCH45 has a circularly permuted double-stranded DNA genome of 212,807 kb and is therefore a jumbophage (i.e. phages with genomes >200kb; Figure 1B)^15^. Sequence analysis of individual genes or the complete genome revealed that PCH45 is highly divergent from known jumbophages, including the well characterised *Pseudomonas Phikzviruses* (Figure 1C)^16,17^. Indeed, its closest relatives, *Erwinia* phage PhiEaH1 and *Serratia* phage 2050HW, showed little sequence conservation (Figure S1). In summary, we identified a unique phage distinct from other described jumbophages.

**Figure 1.**
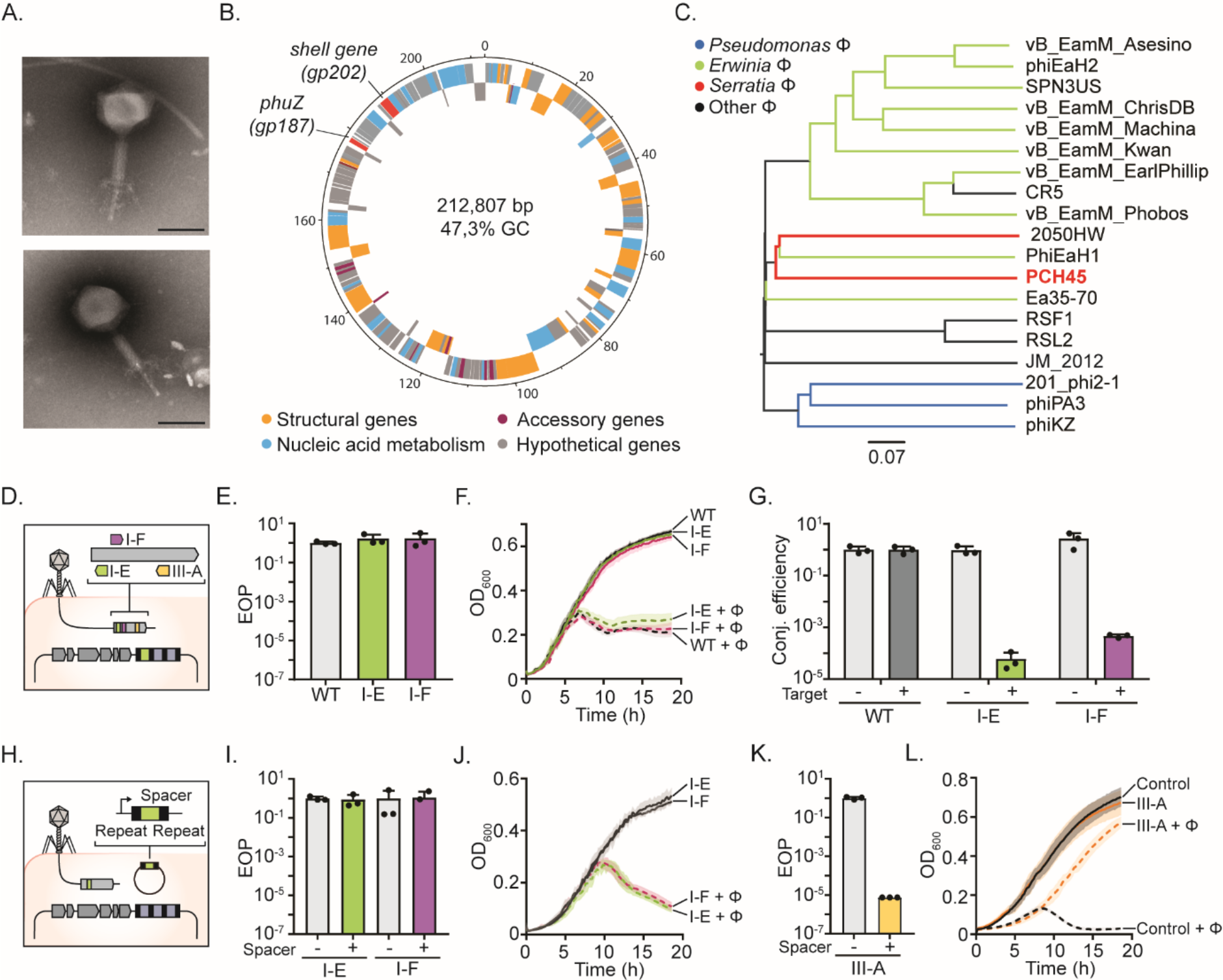
Jumbophage PCH45 evades type I, but not type III, CRISPR-Cas immunity. **A.** Electron micrographs of PCH45 (scale bar: 100 nm) **B.** PCH45 genome; genes on the positive (outer circle) or negative (inner circle) strand are indicated. Gene classification: nucleic acid metabolism (cyan), structural (orange), accessory (purple) and hypothetical (grey). The *phuZ* (*gp187*) and shell (*gp202*) genes are red. **C.** Phylogenomic Genome BLAST Distance Phylogeny tree of jumbophages. Branch lengths are scaled according to the distance formula (*d*_6_). **D.** Major capsid gene (*gp033*) targets of chromosomal type I-E (S1), I-F (S2) and III-A (S3) spacers. Phage resistance by **E.** efficiency of plaquing (EOP) or **F.** plate reader assays for strains with no (WT; control), type I-E (S1, PCF592) or I-F (S2, PCF574) spacers. **G.** Conjugation efficiency of untargeted (pPF1123) and targeted (*gp033*, pPF1443) plasmids into strains in D and E. **H.** Schematic of spacers expressed from mini-CRISPR arrays. Phage resistance by **I.** EOP or **J.** plate reader assays for strains with a type I-E (S1, pPF1459) or I-F (S2, pPF1462) spacer in mini-CRISPR arrays. Plasmids lacking spacers (I-E: pPF974 and I-F: pPF975) were controls. Phage resistance by **K.** EOP or **L.** plate reader assays for strains with a type III-A (S3, pPF1467) spacer in a mini-CRISPR array. A mini-CRISPR array lacking spacers (pPF975) was the control. Data are presented as mean ± SD (n=3). In E, I and K moi=0.001.

To test if the *Serratia* type I CRISPR-Cas systems elicit jumbophage resistance, we generated strains with chromosomal anti-*gp033* (major capsid) spacers in CRISPR1 (I-E) and CRISPR2 (I-F) (Figure 1D, S2A and Table S4). These anti-PCH45 spacers failed to provide jumbophage resistance on plate or liquid cultures (Figure 1E&F, S2B), despite interfering strongly with plasmids (10^5^ fold reduction in conjugation) (Figure 1G). Importantly, these *Serratia* type I systems provide resistance against other phages, including *Siphovirus* JS26^18^. The jumbophage appeared to avoid type I immunity in an anti-CRISPR-independent manner, since no known *acr* genes were detected in the PCH45 genome. Furthermore, the jumbophage DNA was sensitive to digestion by restriction enzymes and no genes encoding known DNA modification enzymes were detected in the genome, indicating that DNA modification was not obstructing CRISPR-Cas defence (Figure S2E). It was also possible that CRISPR-Cas expression was insufficient in *Serratia* to provide resistance; however, the jumbophage still evaded type I immunity when crRNAs were overexpressed from mini-CRISPR arrays (Figure 1H-J and S2C-D). We next examined type III immunity by expressing a spacer targeting the jumbophage major capsid mRNA. In contrast to the type I systems, the type III-A system provided robust phage resistance (Figure 1K&L). Thus, the type III-A system protected against the *Serratia* jumbophage, whereas the type I systems were evaded in an unknown process.

We were interested in how the jumbophage evaded type I, yet was susceptible to type III immunity. Recently, three *Pseudomonas Phikzvirus* phages were shown to produce nucleus-like structures during infection^6,7^. The phage nucleus is surrounded by a shell of phage proteins and is positioned in the cell centre by a phage-encoded tubulin spindle (PhuZ)^7,19,20^. The *Serratia* jumbophage, despite bearing little similarity to the *Phikzvirus* genus, encodes a tubulin homologue (*gp187*) and a potential shell protein (*gp202*) (Figure 1B). These proteins have low sequence identity (16.5% and 19.9% at the amino acid level for PhuZ and the shell protein, respectively) to those in the *Pseudomonas* phage phiKZ (type species of the *Phikzvirus* genus) (Figure S3A&B). Therefore, we hypothesised the *Serratia* jumbophage produces a nucleus-like compartment upon infection. *Serratia* was infected with PCH45 and confocal microscopy was used to visualise DNA and membranes. During infection we observed circular DNA foci, consistent with nucleus-like structures (Figure 2A). By contrast, DNA was evenly distributed in uninfected controls. Thirty minutes after infection, most phage nuclei were either localised centrally (n=102; 61%), or towards the cell poles (n=64; 39%). To test if the DNA foci were encapsulated by a phage shell, the putative shell protein was tagged (*mEGFP*-*gp202*). Upon phage infection, the tagged shell protein assembled into a spherical structure enclosing the phage DNA but no shell was formed without infection (Figure 2B). Therefore, phage infection leads to DNA accumulation within a phage-encoded protein shell that includes protein Gp202.

**Figure 2.**
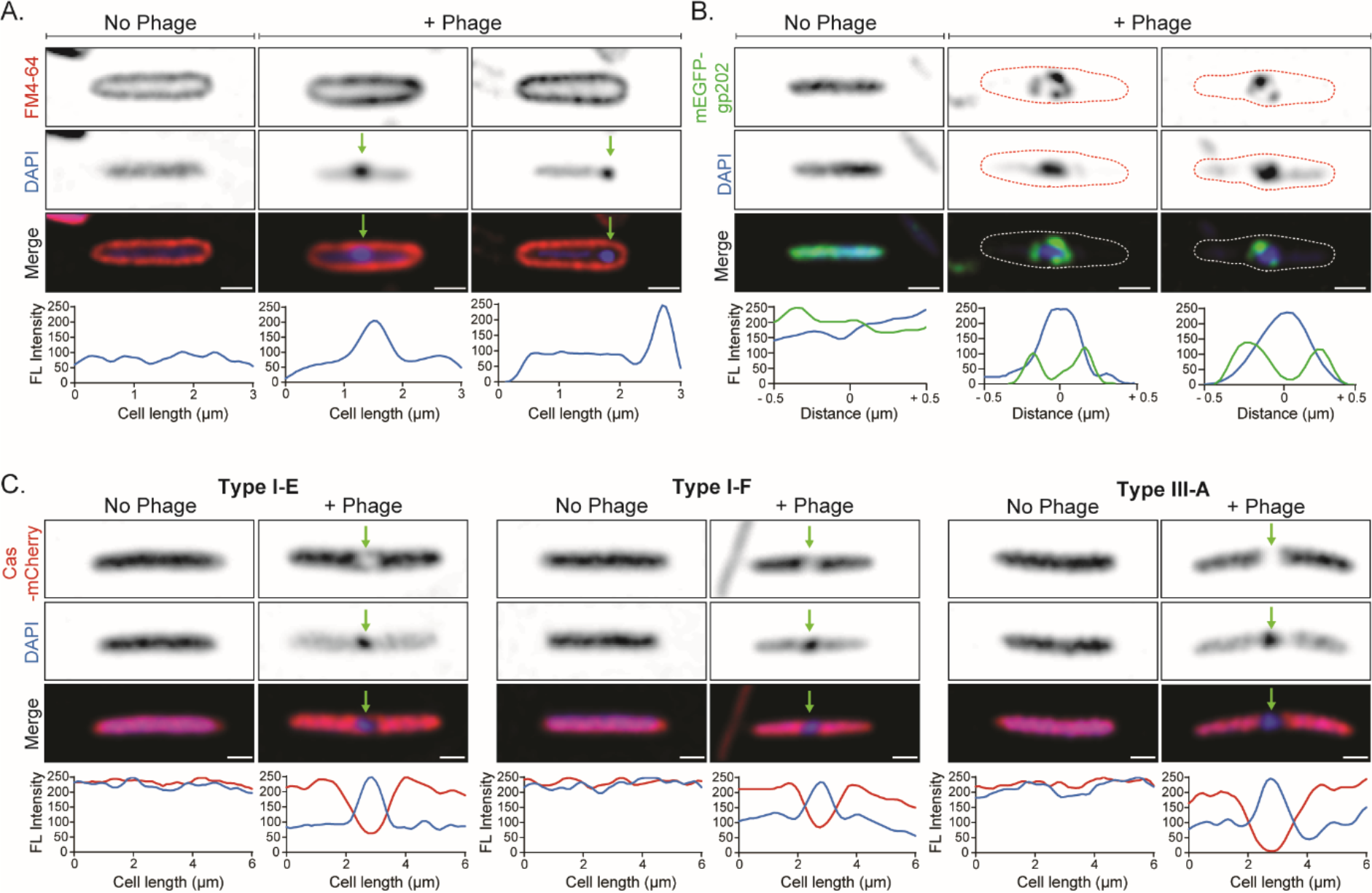
The jumbophage assembles a DNA-containing protein shell during infection that excludes CRISPR-Cas complexes. **A.** Nucleus-like structures (arrows) form in *Serratia* WT infected with PCH45 (+ Phage). Uninfected cells (No Phage) do not form DNA foci. DNA (blue) and membranes (red) were stained with DAPI and FM4-64, respectively. Quantifications show the fluorescence (FL) intensity distribution of DAPI along the cell length of the representative single cells. Scale bar: 1 μm. **B.** Gp202 forms a shell (green) around the DNA foci (blue). Quantifications show the fluorescence (FL) intensity profile of the DNA and shell in the representative single cells. The dashed lines outline the cell shape. Scale bar: 1 μm. **C.** Cas type I-E, I-F and III-A complexes (red) are excluded from DNA (blue) in infected cells (+ Phage) (arrows). Scale bars: 1 μm. Graphs show the fluorescence (FL) intensity distributions of the Cas complexes (red) and DNA (blue) across the cell length of representative single cells. All images were collected 30 min post infection (moi=8).

In *Pseudomonas* Phikzvirus 201 Φ2-1, shell formation allows the selective translocation of proteins into the phage nucleus, restricting other proteins to the cytoplasm^6^. We hypothesised that the *Serratia* jumbophage evades DNA targeting due to the nucleus-like compartment excluding Cas proteins from phage DNA. Therefore, we monitored CRISPR-Cas interference complex localisation during jumbophage infection with the large subunit of all systems tagged with mCherry (I-E, *cas8e*; I-F, *cas8f*; and III-A, *cas10*). All tagged systems retained interference activity against the CRISPR-sensitive phage JS26 (Figure S3C). For all CRISPR-Cas types, the interference complexes were localised in the cytoplasm external to the phage nucleus (Figure 2C). Together, this shows that the CRISPR-Cas complexes are spatially excluded from the phage nucleus, preventing access to the phage DNA.

Replication and transcription of *Phikzvirus* DNA occurs inside the nucleus-like compartment and mRNA is transported to the cytoplasm for translation^6^. This is consistent with the *Serratia* jumbophage being protected from type I systems (target DNA), while remaining sensitive to type III (targets both RNA and DNA)^21,22^. To further investigate the role of type III RNA targeting in jumbophage defence, we tested a panel of crRNAs that target different PCH45 genes (Figure 3A). In agreement with RNA targeting, all anti-sense crRNAs inhibited jumbophage infection in plate and liquid assays, whereas crRNAs sense to transcribed mRNAs provided no protection (Figure 3B-D). Targeting was unaffected by whether the target RNA was predicted to be expressed in an early, middle or late stage of infection (Figure 3A&B).

**Figure 3.**
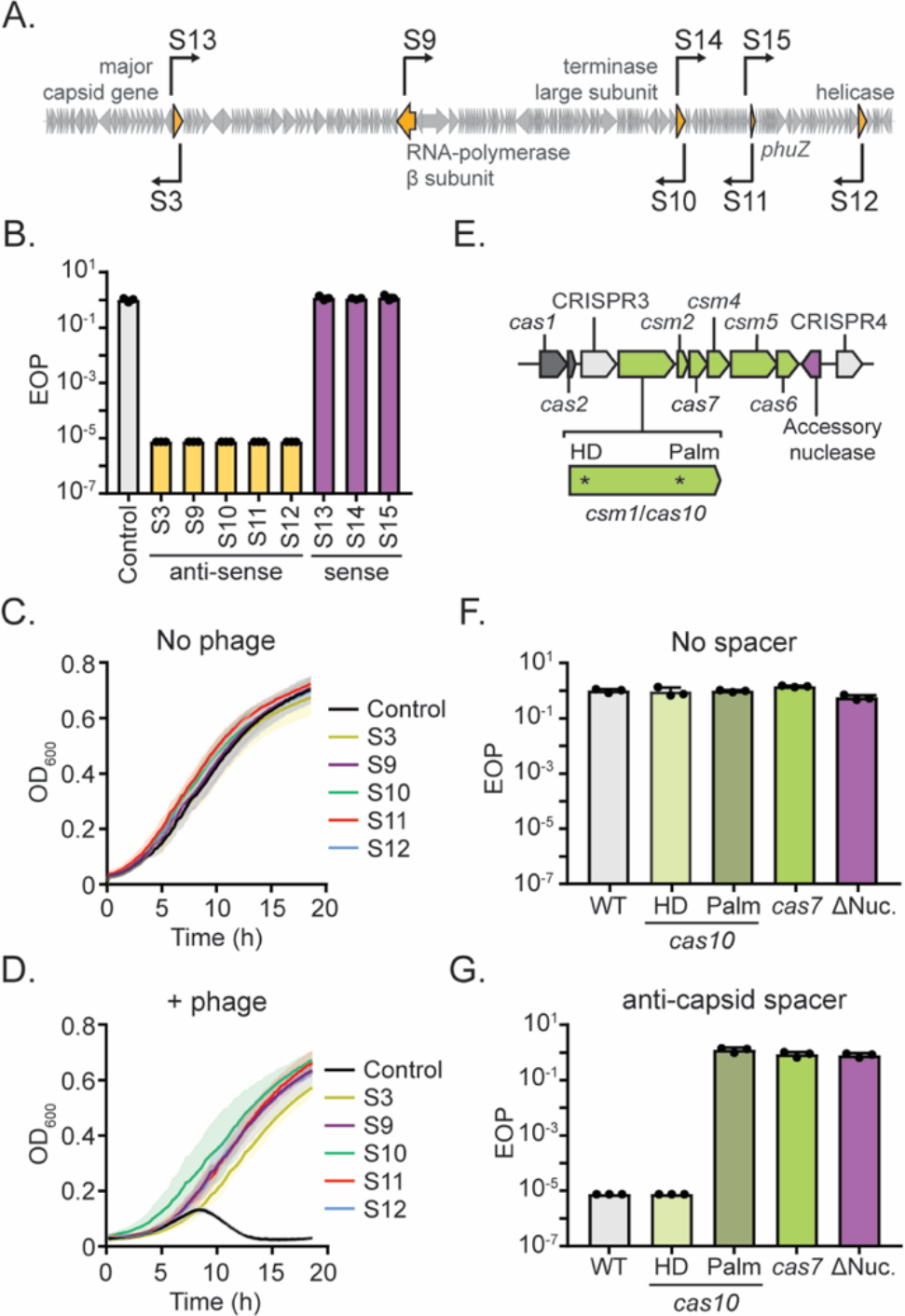
Jumbophage infection is inhibited by the type III CRISPR-Cas system. **A.** PCH45 phage genome indicating targets of the type III-A spacers: major capsid (*gp033*; S3 and S13), RNA-polymerase beta subunit (*gp084*; S9), terminase large subunit (*gp159*; S10 and S14), tubulin-like protein (*gp187*; S11 and S15) and helicase (*gp217*; S12). **B.** EOP assay for strains expressing type III-A spacers from mini-CRISPR arrays (Control; pPF975, S3; pPF1467, S9; pPF1466, S10; pPF1469, S11; pPF1470 and S12; pPF1468). Plate reader assay of strains expressing the spacers in B. either **C.** uninfected or **D.** infected with PCH45 (moi=0.001). **E.** *Serratia* type III-A locus: adaptation genes (grey), CRISPR arrays (light grey), interference genes (green) and accessory nuclease (pink) are shown. EOP assay with **F.** no spacer (pPF975) or **G.** with an anti-capsid spacer (S3) expressed from a mini-CRISPR array in the WT or type III-A mutants: *cas10H17A, N18A* (HD), *cas10D618A, D619A* (Palm), *cas7D34A*, and the nuclease deletion. Data are presented as mean ± SD (n=3).

In type III-A systems, RNAs are recognised by the Csm effector complex^21,22^, triggering sequence-specific RNA cleavage by Cas7^23^. In addition, the nuclease (HD) domain in Cas10 promotes non-specific DNA cleavage^24^. RNA binding also activates the Cas10 Palm domain, which synthesises cyclic oligoadenylate secondary messengers that activate non-specific RNases, causing collateral RNA degradation^25–27^. The *Serratia* type III-A system has two CRISPR-arrays (CRISPR3 and CRISPR4), an operon encoding the adaptation complex and an operon encoding the Csm complex (Figure 3E). In addition, a hypothetical nuclease is convergently transcribed between *cas6* and CRISPR4. To investigate RNA targeting by the *Serratia* type III-A system in the inhibition of jumbophage infection, catalytic mutants of the key proteins involved in RNA and DNA cleavage were tested with multiple spacers (Figure 3F&G and S4A). As predicted, a Cas10 HD mutation (*cas10*^*H17A, N18A*^) did not affect type III-A immunity, indicating that DNA cleavage is not necessary for jumbophage resistance. In contrast, active site mutations in Cas7 (aka Csm3; *csm3*^*D34A*^)^23^ or the Cas10 Palm domain (*cas10*^*D618A, D619A*^), which disrupts cyclic oligoadenylate signalling^26^, abolished phage resistance. Moreover, resistance was also lost when the accessory nuclease was deleted. The same effects on interference of plasmid conjugation were observed for all mutants and restoration of the WT copy of each mutant gene complemented CRISPR-Cas activity (Figure S4B & S4C). Together, jumbophage immunity requires the RNA-targeting and cyclic oligonucleotide signalling capabilities of the type III-A CRISPR-Cas system.

We hypothesised that type III CRISPR-Cas immunity against nucleus-forming jumbophages would occur in natural environments, whereas type I immunity would be rare. To test this hypothesis, we analysed type I-E, I-F and III spacers from ~160,000 bacterial genomes and identified their targets in isolated phage genomes and viral contigs from global metagenomes (Table S2). For total phages, many targets were identified for all systems (Figure 4A). Consistent with our model, targets of type III spacers were significantly enriched in jumbophages (i.e. >200 kb) (Figure 4A&B) and further enriched in nucleus-forming phages that encode homologues of both shell and tubulin proteins (χ^2^ test <0.001) (Figure 4C). Multiple examples of type III systems targeting nucleus-forming jumbophages were present in diverse classes of proteobacteria. In contrast, type I-E and I-F spacer matches were depleted in jumbophages and those defined as nucleus-forming (Figure 4B&C and S4E). In conclusion, both experimental and bioinformatic data provide evidence that type III CRISPR-Cas immunity against jumbophages is widespread in nature, but that these phages evade type I immunity.

**Figure 4.**
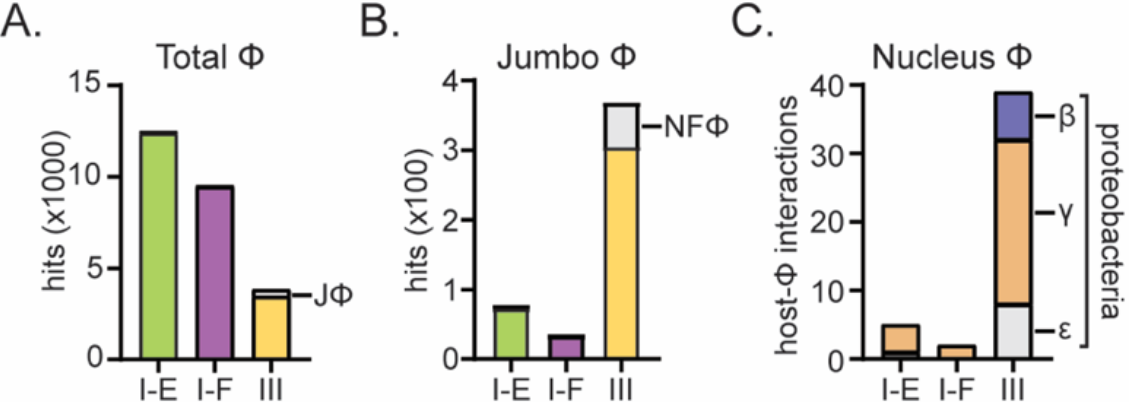
Jumbophages are targeted by the type III CRISPR-Cas systems in different bacterial classes. Number of spacers in type I-E, I-F or type III systems matching **A.** total *Caudovirales* phages with jumbophages (JΦ) in grey, **B.** jumbophages (>200 kb phages) with nucleus-forming phages (NFΦ) in grey) and **C.** unique host-nucleus-forming phage interactions.

We have discovered a jumbophage that evades DNA targeting by two native type I CRISPR-Cas systems while retaining sensitivity to the RNA targeting capabilities of the type III-A system. We propose that resistance is conferred by the formation of a nucleus-like structure in the bacterial cytoplasm that physically shields DNA, but not RNA, from cytoplasmic CRISPR-Cas effector complexes. This concept of exclusion-defence was proposed earlier for *Phikzviruses*^7^ and is supported by a recent unpublished study showing phage DNA, but not RNA, is protected from immune systems different to those tested in this study^28^. Our work on a unique jumbophage that infects a different order of bacteria, and bears little similarity to the *Phikzvirus* genus, coupled with our bioinformatic analyses, provides evidence that the phage nucleus is a widespread counter-defence strategy amongst jumbophages. This manner of immune evasion leads to the prediction that the phage nucleus would provide broad protection from diverse DNA-targeting defence systems. This quality makes nucleus-forming phages prime candidates for phage-based therapies. Importantly, despite DNA protection, RNA export to the cytoplasm is a vulnerability of jumbophages that can be exploited by type III CRISPR-Cas systems. It is likely that jumbophage infection has selected for the observed widespread type III RNA-targeting immunity in strains already possessing DNA-based defences.

## Methods

Detailed Methods are provided in the Supplementary Information.

## Acknowledgements

This work was supported by the Marsden Fund from the Royal Society of New Zealand. LMM was supported by a University of Otago Doctoral Scholarship. We thank staff of the Otago Micro and Nano Imaging facility for assistance with Electron and Confocal Microscopy and the Otago Genomics Facility for genome sequencing. We thank members of the Fineran laboratory for helpful discussions, Saadlee Shehreen, Tom Nicholson and Xochitl Morgan for bioinformatic advice.

## Author contributions

LMM, SLW, CW and LFG performed experiments. LMM, SLW, SAJ and CW generated strains and plasmids. LMM, SLW and LFG performed microscopy. LMM and SAJ performed bioinformatic analysis with input from PPG and PCF. LMM and PCF conceived the project with input from all authors. PCF supervised the project. LMM and PCF wrote the manuscript. All authors edited the manuscript.

## Materials & correspondence

Requests should be made to

