## Supplementary Information for "A nucleus-forming jumbophage evades CRISPR-Cas DNA targeting but is vulnerable to type III RNA-based immunity"

### Methods

#### Bacterial strains, plasmids and culture conditions

The plasmids and primers used in this study are listed in Tables S3 and S4 and bacterial strains are listed in Table S5. Plasmids were all confirmed by sequencing, transformed into *E. coli* ST18 and introduced into *Serratia* strains by conjugation. *Serratia* sp. ATCC 39006 (*Serratia*) and *E. coli* were grown at 30ºC and 37ºC in Lysogeny Broth (LB) under shaking conditions (160 rpm). When grown on plates, containing 1.5% w/v agar in LB (LBA), the strains were incubated at an appropriate temperature until colonies appeared. Media were supplemented with ampicillin (Ap, 100 µg/ml), chloramphenicol (Cm, 25 µg/ml), kanamycin (Km, 50 µg/ml), tetracycline (Tc 1 µg/ml) and 5-aminolevulinic acid (ALA, 50 µg/ml) when required.

#### Jumbophage isolation

Jumbophage PCH45 was isolated from sewage samples extracted from the Tahuna Waste Water Treatment Plant in Dunedin, New Zealand (45°54'16.1"S; 170°31'16.8"E). Enrichment for phages infecting *Serratia* was performed by mixing 100 µl of sewage sample with 5 ml of *Serratia* culture and incubating the mixture overnight at 30ºC shaking at 180 rpm. The phage suspension was separated from the cellular debris by centrifugation (2,000 x g for 20 min at 4ºC), diluted and plated onto *Serratia* lawns. Jumbophages were selected based on their small plaque morphology, picked and used to infect a *Serratia* overnight culture. Serial dilutions of the lysate were plated onto LBA plates and grown overnight. The process was repeated until uniform plaques were obtained to ensure a pure phage stock.

### Preparation of phage stocks and titration

Phage stocks were prepared as described elsewhere[1]. In brief, 100 µl of overnight bacterial culture was mixed with serial dilutions of phage lysate and added to 4 ml of LBA overlay (0.35% w/v) which was then poured onto LBA plates. Plates were then incubated overnight at 30ºC, plates with webbed lysis (almost confluent lysis) had the top agar scraped off and pooled together into a centrifuge tube. A few drops of chloroform ($NaCO_3$ saturated) were added before thoroughly vortexing the mixture to lyse the cells. Finally, a centrifugation step was performed (2,000 x g for 20 min at 4ºC) to separate the virions from the cell debris. The supernatant was placed in a sterile universal for storage and phage titre determined by pipetting 20 µl drops of serial dilutions of the phage stock onto a LBA overlay seeded with 100 µl of *Serratia* overnight culture. Plaques were counted after incubation overnight at 30ºC, with the phage titre represented as pfu/ml. Phage stocks were stored at 4ºC.

### Electron microscopy

To examine the jumbophage by transmission electron microscopy (TEM), 10 µl of high titre phage stock (~$10^9$ pfu/ml) was loaded onto plasma-glowed carbon coated 300 mesh copper grids. After 60 seconds the excess specimen was removed by blotting and 10 µl of 1% w/v phosphotungstic acid (PTA) (pH 7.2), was applied to the grid to stain the samples and blotted off immediately after. The grids were viewed in a Philips CM100 BioTWIN transmission electron microscope (Philips/FEI Corporation, Eindhoven, Holland), and images captured using a MegaView III digital camera (Soft Imaging System GmbH, Münster, Germany).

### Phage DNA extraction and restriction analysis

DNA was extracted from a high titre phage stock (~$10^{10}$ pfu/ml), using the cetyltrimethylammonium bromide (CTAB) method described elsewhere[2]. Briefly, 2.5 ml of phage stock were treated with 100 ng of RNase A and 100 U of DNAse I and incubated at 37ºC for 30 min to clean the phage from bacterial nucleic acid. Enzyme inactivation was performed by the addition of 0.8 ml 0.58 M EDTA (pH 8.0). Next, phage proteins were removed by the addition of proteinase K (20 mg/ml) and incubation at 45ºC for 15 min. CTAB (10% w/v) in 4% NaCl (w/v) was preheated to 55ºC, added to the sample, mixed gently by inversion and cooled on ice for 15 min.

The CTAB-DNA complex was collected by centrifugation (12,000 x g for 15 min) and the pellet was resuspended in 1 ml of 1.2 M NaCl. Phage DNA was precipitated adding 600 µl of isopropanol, washed with 75% ethanol and resuspended in 100 µl of TE buffer. Samples were cleaned with DNeasy Blood & Tissue Kit (QIAGEN) following the manufacturer's instructions. The DNA concentration was determined by fluorometric quantification using Qubit dsDNA HS Assay Kit and the Qubit Flourometer following the manufacturer's instructions.

Restriction digestion was performed on PCH45 DNA to check for DNA modification. Digestion of 1 µg of PCH45 genomic DNA was performed with different restriction enzymes (NEB). The samples were incubated for 2 hours at 37°C and separated on a 1% agarose gel alongside the 1 kb+ ladder (Thermofisher), stained with EtBr and visualized under UV light.

**Genome sequencing, annotation and comparative genomics**

Library preparation and sequencing was performed by Massey Genome Service (Massey University, Palmerston North, New Zealand). First, DNA libraries were prepared with Illumina Nextera XT DNA Library Preparation Kit_V2. The QC was checked using the Quant-iT dsDNA HS Assay for quantification and analysed using SolexaQA++, fastQC and fastQsceen. Sequencing was performed on an Illumina MiSeq (2 x 150 bp), and resulting reads were processed and trimmed using the SolexaQA++ software. The resulting reads were assembled de novo using SPAdes 3.9[3] and annotated with RASTtk[4]. Further annotation was performed manually by analyzing the BLASTp hits for each independent CDS identified by RASTtk. tRNAscan-SE v. 2.0[5] was used to identify putative tRNAs. The genome was visualized using DNAplotter[6]. Similar phage genomes were identified using PAirwise Sequence Comparison (PASC)[7] and pairwise genome comparisons were performed with Easyfig[8]. Finally, the taxonomic trees were built using VIrus Classification and Tree building Online Resource (VICTOR)[9] and modified with FigTree.

**Data availability statement**

The genome sequence of bacteriophage PCH45 was deposited in the GenBank database under accession number MN334766.

### Generation of native type I-E and I-F anti-phage strains

*Serratia* strains harbouring anti-PCH45 spacers in their type I-E and I-F chromosomal arrays were obtained by primed spacer acquisition[10]. Plasmids primed by the type I-E and I-F systems (pPF1255 and pPF1256) were generated as follows. The major capsid gene (*gp033*) was amplified by PCR from phage gDNA using PF2231/PF2232. The insert was digested with SpeI and KpnI and cloned into the priming vectors (pPF1125 and pPF1126) that were previously digested with the same enzymes. pPF1125 and pPF1126 contained a protospacer primed by spacer 1 from either the type I-E and the I-F CRISPR-Cas systems in *Serratia*. Plasmids were transformed into *E. coli* ST18, plated onto LBA + ALA + Km and grown overnight at 37ºC. The vectors were conjugated into *Serratia* by mixing equal volumes of donor and recipient and plating the mating spot onto LBA + ALA. The mating spot was streaked onto LBA + Cm to select for *Serratia* transconjugants. *Serratia* colonies were grown overnight in the absence of antibiotic, to allow plasmid loss. The cultures were passaged for 3 days by inoculating LB with 5 µl of overnight culture. Each day serial dilutions were plated and incubated at 30ºC until colonies appeared on the plate. The clones were screened by PCR in search of CRISPR array expansion (PF1989 – PF1887 for CRISPR1 and PF1990 – PF1889 for CRISPR2). In cases where CRISPR expansion was observed, PCR products were sequenced to determine if the spacers acquired corresponded to the plasmid backbone or the phage fragment (Table S3 and S6). The clones were also patched onto LBA + Cm to check for plasmid loss.

### Plasmid expression of anti-phage spacers

For some experiments, spacers were expressed from plasmid mini-CRISPR arrays. These plasmids were constructed by cloning anti-PCH45 and anti-JS26 spacers into plasmid mini-CRISPR arrays. Briefly, reverse complement primers carrying spacer sequences targeting phage genes flanked by two BsaI restriction sites were annealed (details in Table S3 and S4). The annealed primers were cloned into plasmids pPF974, pPF975[11] and pPF976 (expression plasmids carrying type I-E, I-F and III-A mini-CRISPR arrays) previously digested with BsaI. pPF976 was constructed in the same manner as pPF974/5[11] using primers PF1981 and PF1982. The plasmids carrying the mini-CRISPR arrays were introduced into *E. coli* ST18 by transformation and then conjugated into *Serratia* (and derivative mutants). Expression of crRNAs was induced by addition of IPTG (0.1 mM) and strains were grown in the presence of Km for plasmid maintenance.

**Phage resistance efficiency of plaquing assay**

To assess the infectivity of PCH45 and JS26 on different *Serratia* strains, efficiency of plaquing (EOP) assays were performed. A soft LBA overlay (0.35% w/v) containing 100 µl of bacterial culture was poured onto an LBA plate. Serial 10-fold dilutions of high titre phage stock (~$10^9$ pfu/ml) were spotted (20 µl) onto the agar overlay and plates were incubated overnight at 30ºC. The EOP was calculated as the ratio of pfu/ml produced on tested strains to the pfu/ml on the control *Serratia* strain. All the conditions were repeated in triplicate and plotted as the mean plus or minus the SD.

**Phage resistance infection time courses**

*Serratia* cultures were grown from an initial $OD_{600nm}$=0.05 at 30ºC, shaking (160 rpm) until reaching an $OD_{600nm}$= 0.3 (exponential phase). The cultures were diluted to an $OD_{600nm}$ =0.05, 180 µl aliquots were pipetted into a 96-well plate and 20 µL of diluted phage lysate was added to produce an moi=0.001. The 96-well plate was incubated in a Varioskan Flash plate reader (Thermo Scientific) at 30ºC with 240 rpm shaking and $OD_{600nm}$ measurements measured in 12 min increments for 20 h. Each condition was repeated in triplicate with data plotted as the mean plus or minus the SD. For strains with anti-phage spacers expressed from a plasmid, the media was supplemented with Km and IPTG (0.1 mM).

**Construction of mEGFP/Shell protein expression plasmid**

A plasmid (pPF1956) for expression of an mEGFP-tagged shell protein was generated by PCR amplification of the shell gene (*gp202*) using PF3825/PF3812 and *mEGFP* using PF3826/PF3827 from phage gDNA and the gBLOCK PF3809, respectively. The vector pQE-80L-oriT stuffer was digested with SphI and KpnI and assembled with the two inserts using Gibson Assembly.

### Construction of fluorescently tagged Cas complexes

Cas complexes were fluorescently tagged by fusing *mCherry2* to the N-terminal region of each large subunit (*cas8e*, *cas8f* and *cas10*). Plasmids were constructed by cloning *mCherry2* flanked by the upstream region and the first 500 bp of each gene into the suicide vector pPF1117[11]. The regions were amplified by PCR from gBLOCK PF3810 (*mCherry2*+linker) and *Serratia* WT using primers PF3817/PF3818, PF3811/PF3812, PF3819/PF3820 (*mCherry2-cas8e*, pPF1951), PF3821/PF3822, PF3811/ PF3812, PF3823/ PF3824, (*mCherry2-cas8f,* pPF1953) and PF3813/PF3814, PF3811/PF3812, PF3815/ PF3816 (*mCherry2-cas10,* pPF1955). The inserts were cloned into pPF1117, previously digested with SphI and SalI, using Gibson Assembly. The fluorescently tagged Cas complexes were introduced into the chromosome by homologous recombination. The suicide vectors for *cas8e* (pPF1951), *cas8f* (pPF1951) and *cas10* (pPF1955) were conjugated into *Serratia* and recombination was selected by growth on LBA with 20% w/v sucrose and colonies were screened for Cm sensitivity as described elsewhere[12,13]. This gave strains PCF734 (*mCherry-cas8e*), PCF736 (*mCherry-cas8f*) and PCF732 (*mCherry-cas10*). To optimise the expression of the fluorescent Cas complexes, an IPTG-inducible T5 promoter was inserted upstream each complex. *mCherry2* was amplified by PCR from gBLOCK PF3810 using primers PF4005/PF4007 and cloned into pPF1813[14] previously digested with EcoRI and XmaI, using Gibson Assembly to yield plasmid pPF2036. Plasmid pPF2036 was inserted at the *mCherry2* site by homologous recombination in PCF732, PCF734, PCF736 and the function of the complexes was confirmed in EOP assays using target plasmids pPF1473, pPF1485 and pPF1489 against *Serratia* Siphovirus JS26 as described earlier.

### Confocal microscopy

To observe infected and uninfected bacteria utilizing microscopy, cultures of bacteria were grown overnight from single colonies, these were then sub-cultured and grown to an $OD_{600}$ between 0.2 to 0.3 (incubated at 30ºC with 180 rpm shaking). For infected samples, phage PCH45 was added to an MOI of ~8 with an infection time of 30 min. Aliquots of 3 ml of each culture were then pelleted and washed with minimal medium (40 mM $K_2HPO_4$, 14.7 mM $KH_2PO_4$, 0.1 % $(NH_4)_2SO_4$, 0.4 mM $MgSO_4$ and 0.2 % sucrose, pH 7.1) before being stained with DAPI (4 µg/ml) and FM4-64 (12 µg/ml) for 20 to 30 min. Stained samples were then washed twice with minimal medium before being resuspended in 100 µl of minimal medium. To suspend samples, 15 µl aliquots of each sample were mixed with 15 µl of molten 1.2% agarose (made up in minimal media) before being sealed onto microscope slides with a cover slip.

To observe the formation of a shell structure, a tagged putative shell gene (*mEGFP-gp202*) was expressed from plasmid pPF1956. Due to leaky expression of the *mEGFP-gp202* shell gene, no induction was required. To observe the localization of interference complexes upon phage infection type I-E, type I-F and type III-A complexes were tagged with mCherry2 as described earlier. Expression of mCherry2-tagged type I-E, I-F and III-A Cas complexes was induced with the addition of IPTG (10 µM) at the time of infection.

Images were acquired using a CFI Plan APO Lambda 100 X 1.49 NA oil objective (Nikon) on the multimodal Imaging Platform Dragonfly (Andor Technology, Oxford Instruments) equipped with 405, 488, 561 and 637 nm lasers built on a Nikon Ti2-E microscope body with Perfect Focus System (Nikon Instruments, Japan). Data were collected in Spinning Disk 40 µm pinhole mode on the iXon888 EMCCD camera with 2X optical magnification using Fusion v1.4 software. Z stacks were collected with 0.1 µm increments in the z axis using an ASI stage with 500 µm piezo z-drive. Data were visualised using Fiji software. Microscopic images were further processed by the deconvolution algorithm in the Huygens Scientific Volume Imagine (SVI) Image Analysis Program.

**Image Analysis**

For quantification of the fluorescence intensity distribution in single cells in Figure 2, the length of cells and the length of nucleoids were traced using Fiji software. Subsequently, the intensity profile along the linescan were measured and plotted in Prism Graphpad.

**Construction of chromosomal type III-A mutants**

To mutate *cas7* and the accessory nuclease in the type III-A system, these genes were first replaced in the chromosome with a Km cassette by homologous recombination. To generate the knock out constructs (pPF1929; *cas7*, pPF1932; nuclease), the up/downstream regions of each gene were amplified by PCR using primer pairs PF3750/PF3585, PF3754/PF3751 (*cas7*) and PF3743/PF3745, PF3748/PF3749 (nuclease) using *Serratia* DNA as a template. The Km cassette was cloned from pSEVA211[15] using primer pairs PF3752/PF3753 (for pPF1929) and PF3746/PF3747 (for pPF1932). The suicide vector pPF1117[11] was digested with SphI and SalI and the inserts were cloned using Gibson Assembly (HiFi DNA Assembly Master Mix, NEB). To generate the *cas10* knock out vector (pPF927), primer pairs PF1934 (SalI site)/PF1935 (BamHI site) and PF1936 (BamHI site)/PF1937 (SphI) were used to amplify by PCR the *cas10* up/downstream regions from *Serratia* WT colonies as DNA template. The two inserts were cloned with a three-part ligation including suicide vector pPF923[11] previously digested with SalI and SphI. The $Km^R$ marked deletion strains were generated (PCF682; *cas7* and PCF685; nuclease) using plasmids pPF1929 and pPF1932 via homologous recombination as described for the tagged Cas complex strains. A markerless *cas10* deletion (PCF303) was constructed by homologous recombination using plasmid pPF927.

Plasmids for site-directed mutagenesis of *cas10* were constructed as described below. Point mutations were introduced by overlap extension PCR from *Serratia* DNA template using primers carrying the altered sequence and a complementary region with an overlapping primer. Amplification was performed using the following primer pairs: PF3756/PF2167 and PF2166/PF3757 for pPF1936 (Cas10 HD domain mutant); PF3756/PF2127 and PF2126/PF3757 for pPF1938 (Cas10 Palm domain mutant) and PF3756/PF2167, PF2166/PF2127, PF2126/PF2757 for pPF1937 (HD and Palm domain double mutant). For pPF1931 ($cas7^{D34A}$), the up/downstream regions of *cas7* were cloned using primers pairs PF3750/PF3585 and PF3755/PF3751 respectively and *Serratia* WT colonies as DNA template; and primer pairs PF3589/PF3590 (PF3755) and gBLOCK PF3591 as DNA template. To delete the accessory nuclease, pPF1933, was constructed using primer pairs PF3743 (overlap with pPF1117)/PF3745 (overlap with PF3749), PF3749 (overlap with PF3745)/PF3744 (overlap with pPF1117) and *Serratia* as template DNA. All inserts were cloned into pPF1117, previously digested with SalI and SphI, using Gibson assembly.

Type III-A mutant strains PCF683 (*cas7*$^{D34A}$), PCF690 (*cas10*$^{H17A, N18A}$, HD mutant), PCF691 (*cas10*$^{D618A, D619A}$, Palm mutant) the HD/Palm double mutant PCF689 (*cas10*$^{H17A, N18A, D618A, D619A}$) and PCF686 (unmarked deletion of the type III-A accessory nuclease; CWC46_RS19930) were generated by homologous recombination as described for the tagged Cas complex strains. The genes carrying the appropriate point mutations were introduced by homologous recombination with plasmids (pPF1930, pPF1933, pPF1934 and pPF1935) into the *cas7* (PCF682) and *cas10* (PCF303) deletion mutants. To generate the unmarked nuclease mutant (PCF685) pPF1933 was recombined into the marked nuclease mutant (PCF686).

**Complementation of type III-A chromosomal mutants**

The type III-A mutants were complemented by reinserting a WT copy of each of the genes into the mutant backgrounds. Constructs pPF1931 (*cas7*), pPF1934 (accessory nuclease) and pPF1935 (*cas10*) were generated using primers PF3750/PF3751, PF3743/PF3744, PF3756/PF3757 and cloned into pPF1117 as described above. Unmarked genes were reinserted by homologous recombination into strains PCF682 (Δ*cas7*), PCF685 (Δnuclease) and PCF303 (Δ*cas10*) with plasmids carrying WT genes (pPF1931; *cas7*, pPF1934; nuclease and pPF1935; *cas10*).

**Type I-E and I-F plasmid interference assay**

To generate a vector targeted by the anti-PCH45 type I-E and I-F spacers (pPF1443), the priming protospacer in pPF1255 was removed. After digestion with SpeI and SphI, pPF1255 was gel extracted, treated with Mung Bean Nuclease and re-ligated using T4 ligase. The interference ability of *Serratia* strains (PCF592 and PCF547) carrying type I-E and I-F anti-PCH45 spacers was tested in conjugation efficiency assays. Cultures of donor *E. coli* ST18 carrying plasmids pPF1123 (untargeted) or pPF1443 (targeted), and the *Serratia* recipient strains were grown overnight. Cultures were adjusted to an $OD_{600}$=1 and donor and recipient strains were mixed in a 1:1 ratio. The mixture was spotted onto LBA + ALA and incubated overnight at 30°C. Next, the mating spots were scraped from the plate and resuspended in 1 ml PBS. Ten-fold serial dilutions were performed and 10 μl of each were spotted onto LBA (total recipient count) and LBA + Cm (transconjugants).

Conjugation efficiency was calculated as transconjugants (cfu/ml)/ total recipients (cfu/ml). For the type III-A mutants (PCF683, PCF686, PCF689, PCF690, PCF691) and their complemented controls (PCF684, PCF687, PCF688), plasmids pPF781 (untargeted) and pPF1043 (targeted) were used. Conjugation efficiency was performed as described for the type I interference assay, but with the following differences. LBA plates for the mating spots included glucose (0.2% w/v) and arabinose (0.02% w/v) was included in all plates for transconjugant and total recipient counts to induce protospacer (target site) transcription.

**Identification of phages targeted by type III spacers**

Type III, I-E and I-F CRISPR-Cas hosts were identified by the presence of *cas* gene annotations in the RefSeq 95 bacterial genomes database. CRISPR loci were extracted using CRISPRDetect[16] with a cutoff score of 2.5. For predicted type III systems, we excluded CRISPRs with repeats that matched known non-type III systems[17] or with repeat lengths $< 30$ nt or $> 50$ nt, and all spacers $< 25$ nt or $> 45$ nt. For type I-E systems, repeats $< 28$ nt or $> 32$ nt, and spacers $< 28$ nt or $> 34$ nt were excluded. For type I-F systems, repeats $< 26$ nt or $> 30$ nt, and spacers $< 28$ nt or $> 34$ nt were excluded. We searched for matches to the spacers against *Caudovirales* genomes in GenBank and non-eukaryotic viral contigs in the IMG/VR database (July 2018)[18] using GASSST[19] (seed=8, sensitivity=3, match=80, gaps=0). Spacer-target matches were scored along their full-length alignment as +1 match and -1 mismatch. The dinucleotide shuffled control datasets, used to determine appropriate scoring cutoffs for the spacer-target matches (Figure S4A-D), were generated using fasta-shuffle-letters from the MEME Suite[20]. Redundant spacer-target matches, due to similar hosts CRISPRs or phage genomes sequences, were first filtered by selecting the highest scoring match for each unique spacer sequence then merging any remaining redundant host-target matches, such that each spacer is represented only once in the dataset.

**Classification of phages as nucleoid-forming**

Using the PCH45 shell (*gp202*) and tubulin (*gp187*) protein sequences as queries, we identified homologous proteins in GenBank *Caudovirales* genomes ≥ 150kb via iterative HMM searches using jackhmmer[21]. Phages containing homologues to both the shell and tubulin proteins (e-values $< 10^{-10}$ for both) were classified as nucleoid-forming. We then manually curated shell and tubulin protein alignments (MUSCLE)[22] for only the shell and tubulin homologs occurring in nucleoid-forming phages (Figure S4A). Using these alignments we generated HMMs using HMMER3[23] and used the HMMs to classify target phage genomes and IMG/VR contigs as nucleoid-forming (matches to both the shell and tubulin HMMs with e-values $<10^{-6}$). Since many IMG/VR contigs represent incomplete phage genomes belonging to defined viral families (clusters), we also classified contigs (~10% of the nucleoid-forming matches) as belonging to nucleoid-forming phages if there were other contigs within the viral cluster that encoded homologues of both the shell and tubulin proteins.

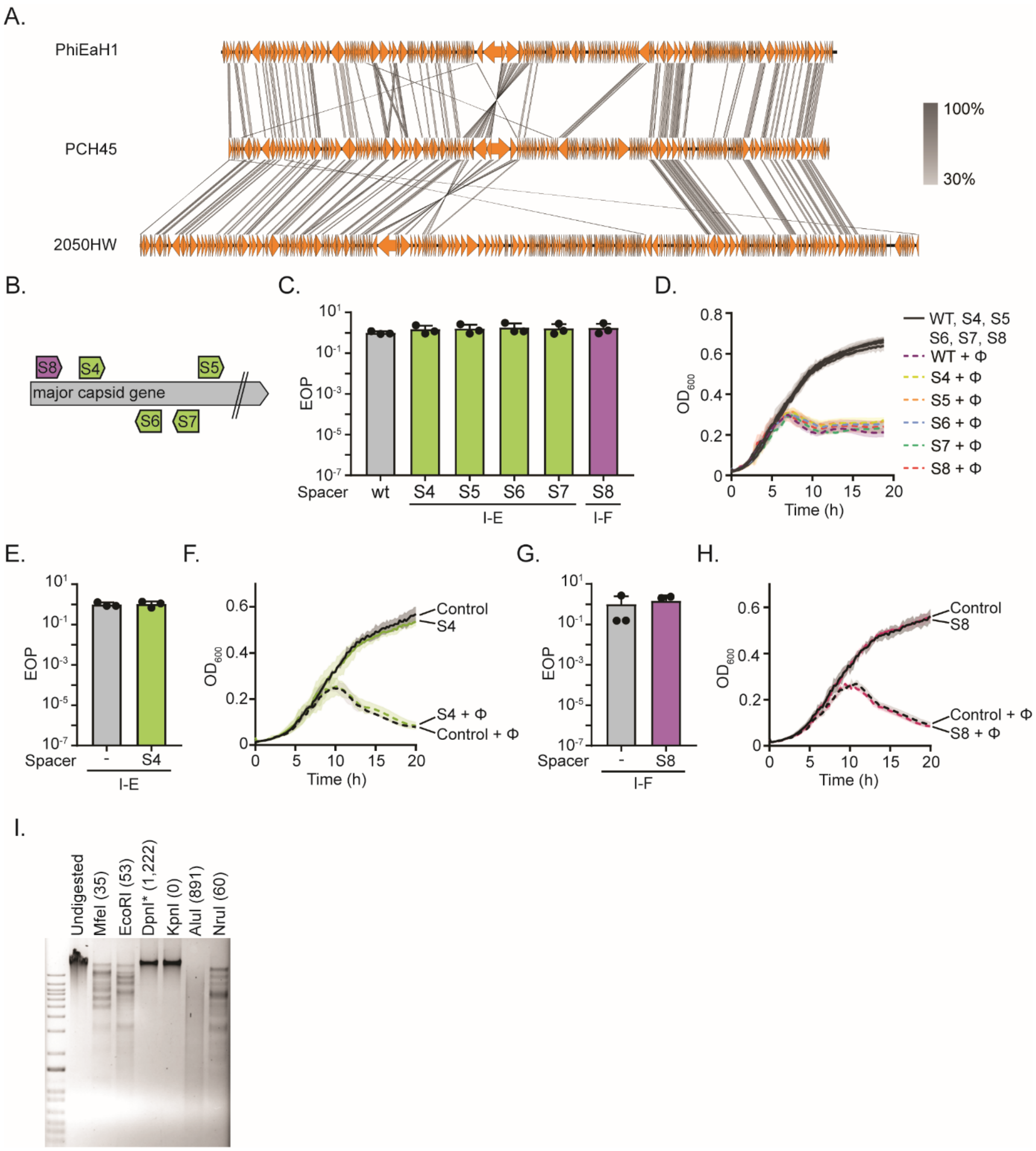


**Figure S1. The jumbophage is resistant to type I CRISPR-Cas immunity A.** tblastx alignment of PCH45 with phages PhiEaH1 and 2050HW. **B.** Target location of chromosomally expressed anti-PCH45 type I-E (S4-7) and type I-F (S8) spacers targeting major capsid gene (*gp033*). Phage resistance measured by **C.** EOP or **D.** plate reader assays for *Serratia* strains with type I-E, (S4, PCF591; S5, PCF593; S6, PCF545; S7, PCF544) and type I-F (S8, PCF548) infected with PCH45. Phage resistance measured by **E.** EOP or **F.** plate reader assays for *Serratia* carrying a type I-E (S4, pPF1460) spacer in mini CRISPR-arrays, infected with PCH45. Phage resistance measured by **G.** EOP or **H.** plate reader assays for *Serratia* carrying a type I-F (S8, pPF1461) spacer in a plasmid borne mini array, infected with PCH45. In C, E and G moi=0.001. **I.** Restriction length fragment polymorphism (RLFP) analysis of phage gDNA treated with restriction enzymes MfeI, EcoRI, DpnI*, KpnI, AluI and NruI. Undigested PCH45 gDNA was used as a negative control.. In parenthesis the number of restriction sites found in the genome of PCH45. (*): Cleaves only when the recognition motif is methylated.

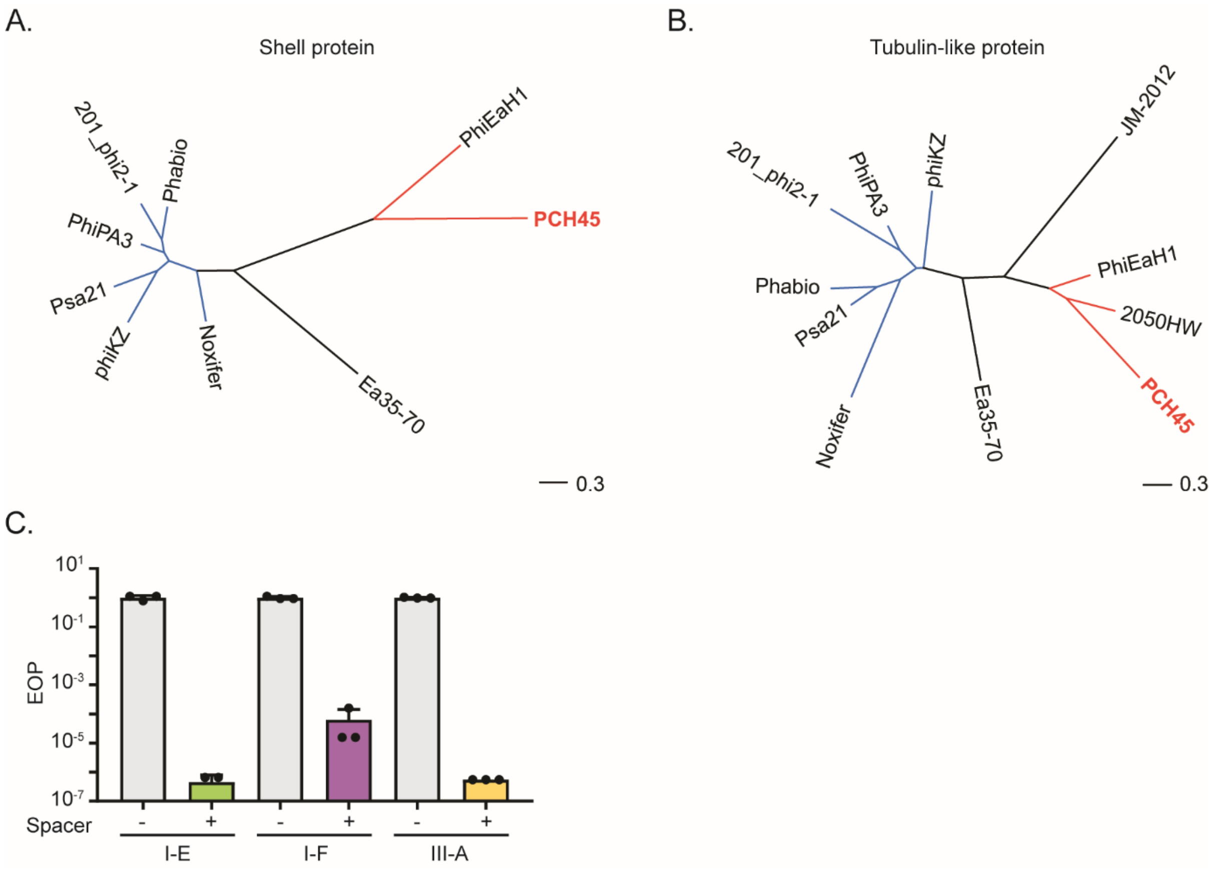


**Figure S2. The shell and tubulin proteins in *Serratia* possess low sequence to homologues encoded by other jumbophages.** Phylogenetic tree of **A.** the shell protein and **B.** PhuZ protein encoded by jumbophages. The tree was generated using ClustaW. The scale bar represents the approximate number of changes per amino acid position. **C.** EOP assay for *Serratia* strains PCF761 (*mCherry2-cse*), PCF763 (*mCherry2-cas8f)* and PCF765 (*mCherry2-cas10*) carrying type I-E, I-F and III-A anti-JS26 spacers in plasmids (pPF1485, pPF1489 and pPF1473 respectively).

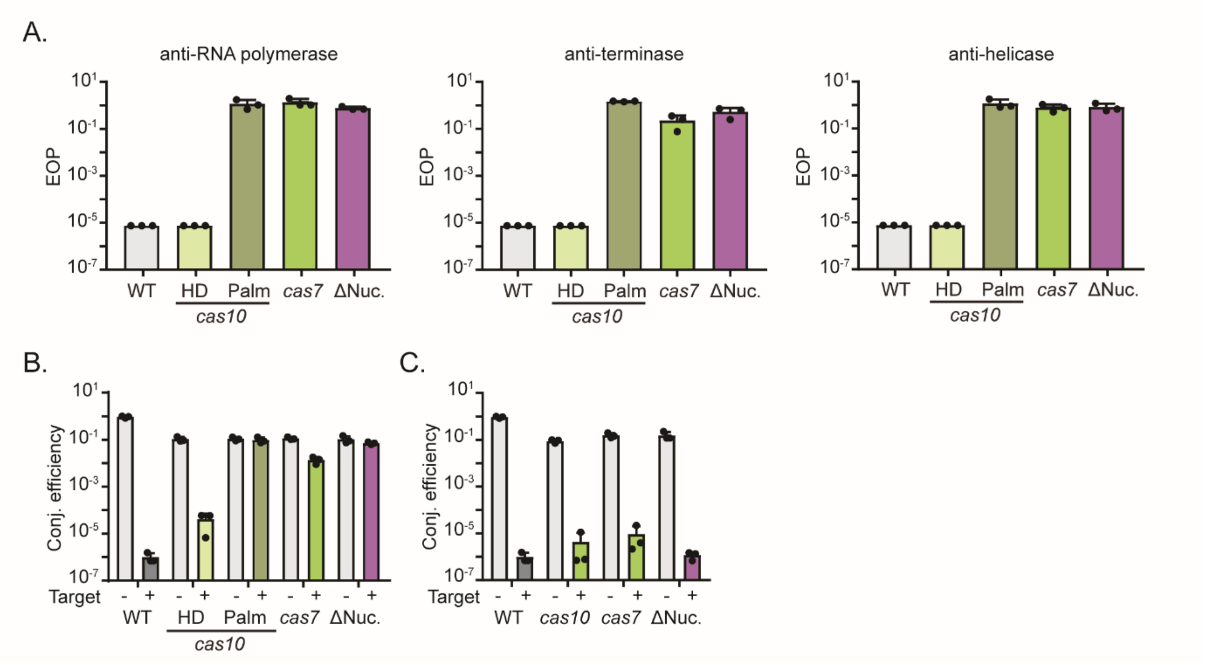


**Figure S3. Type III RNA targeting provides protection against jumbophage infection. A.** EOP assay for type III-A mutant strains: *cas10*$^{H17A,\ N18A}$ (HD domain), *cas10*$^{D618A,\ D619A}$ (Palm domain), *cas7*$^{D34A}$, and the accessory nuclease knock out carrying an anti-PCH45 spacers (RNA polymerase beta subunit, S9; anti-terminase S10; and anti-helicase, S12) overexpressed *in trans* from a plasmid borne mini-array. **B.** Conjugation efficiency assay (cfu/ml) of plasmids pPF781 (naïve) and pPF1043 targeted by the type III-A CRISPR-Cas systems for *Serratia* strains. The type III-A mutants: PCF683 (*cas7*$^{D34A}$), PCF690 (*cas10* HD mutant), PCF691 (*cas10* Palm mutant), PCF689 (*cas10* HD and Palm mutant), PCF686 (Δ accessory nuclease), and **C.** the chromosomal complementation PCF684 (*cas7* wt), PCF688 (*cas10*) and PCF687 (accessory nuclease wt).

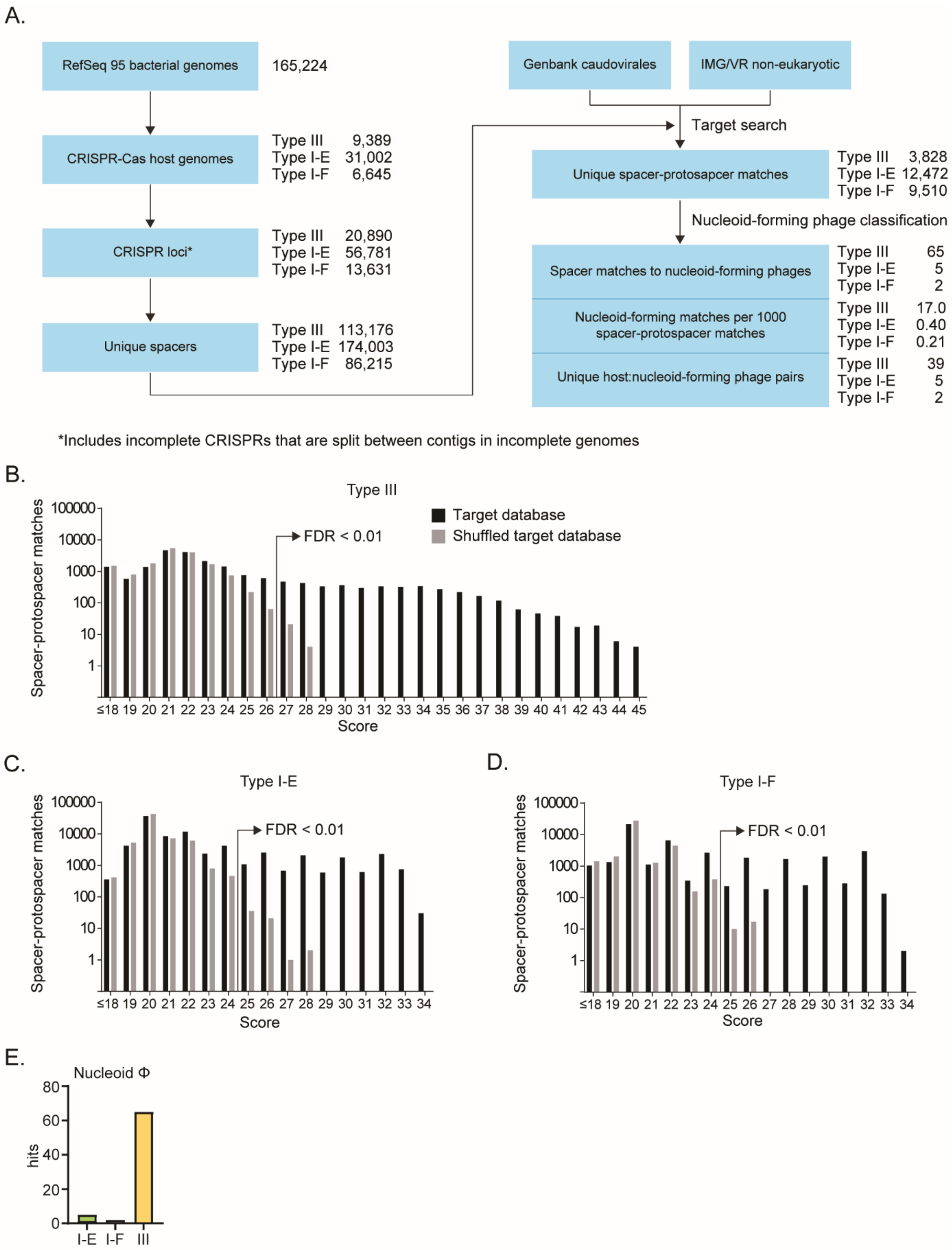


**Figure S4. Type III CRISPR arrays are enriched in jumbophage-targeting spacers. A.** Workflow used to obtain spacer-phage hits. Scores for spacer-target matches for targeted (black) and shuffled (grey) databases for **B.** type III **C.** type I-E and **D.** type I-F CRISPR-Cas systems. Scores with a false discovery rate (FDR) < 0.01 were used as a cut-off to determine the spacer-protospacer hits. **E.** Number of unique spacers in type I-E, I-F or type III systems matching nucleoid forming phages.

**Table S1.** PCH45 genome annotation

| PCH45 ORF name | Protein function | Start (bp) | Stop (bp) | Length (bp) | Strand |
|---|---|---|---|---|---|
| PCH45_001 | Hypothetical protein | 32 | 874 | 843 | + |
| PCH45_002 | Hypothetical protein | 874 | 1290 | 417 | + |
| PCH45_003 | Putative DNA-directed RNA polymerase beta subunit | 1291 | 2658 | 1368 | + |
| PCH45_004 | Structural protein | 2666 | 3310 | 645 | + |
| PCH45_005 | Hypothetical protein | 3398 | 3691 | 294 | + |
| PCH45_006 | Hypothetical protein | 3699 | 4160 | 462 | + |
| PCH45_007 | Hypothetical protein | 4144 | 4272 | 129 | + |
| PCH45_008 | Structural protein | 4339 | 4992 | 654 | + |
| PCH45_009 | Hypothetical protein | 5023 | 5694 | 672 | + |
| PCH45_010 | Structural protein | 6239 | 5757 | 483 | - |
| PCH45_011 | Hypothetical protein | 6664 | 6254 | 411 | - |
| PCH45_012 | Succinate-semialdehyde dehydrogenase | 6858 | 6661 | 198 | - |
| PCH45_013 | DNA polymerase | 8663 | 6858 | 1806 | - |
| PCH45_014 | Structural protein | 8792 | 10075 | 1284 | + |
| PCH45_015 | Hypothetical protein | 10085 | 10549 | 465 | + |
| PCH45_016 | Hypothetical protein | 10552 | 11598 | 1047 | + |
| PCH45_017 | Hypothetical protein | 11595 | 12977 | 1383 | + |
| PCH45_018 | Structural protein | 15963 | 13021 | 2943 | - |
| PCH45_019 | Structural protein | 17146 | 15956 | 1191 | - |
| PCH45_020 | Structural protein | 17175 | 18314 | 1140 | + |
| PCH45_021 | Structural protein | 18324 | 19235 | 912 | + |
| PCH45_022 | Structural protein | 19232 | 19759 | 528 | + |
| PCH45_023 | Hypothetical protein | 19784 | 21193 | 1410 | + |
| PCH45_024 | Hypothetical protein | 21282 | 22613 | 1332 | + |
| PCH45_025 | Hypothetical protein | 22736 | 23758 | 1023 | + |
| PCH45_026 | Phage protein | 23839 | 25392 | 1554 | + |
| PCH45_027 | Structure protein | 25398 | 26786 | 1389 | + |

| PCH45 ORF name | Protein function | Start (bp) | Stop (bp) | Length (bp) | Strand |
|---|---|---|---|---|---|
| PCH45_028 | Hypothetical protein | 26795 | 27385 | 591 | + |
| PCH45_029 | Structural protein | 27385 | 28734 | 1350 | + |
| PCH45_030 | Hypothetical protein | 28727 | 29620 | 894 | + |
| PCH45_031 | Putative helicase | 31232 | 29685 | 1548 | - |
| PCH45_032 | Hypothetical protein | 31282 | 31860 | 579 | + |
| PCH45_033 | Major capsid protein | 31949 | 34282 | 2334 | + |
| PCH45_034 | Structural protein | 34531 | 35193 | 663 | + |
| PCH45_035 | Hypothetical protein | 35301 | 35714 | 414 | + |
| PCH45_036 | Hypothetical protein | 35714 | 36157 | 444 | + |
| PCH45_037 | Hypothetical protein | 36173 | 36505 | 333 | + |
| PCH45_038 | Hypothetical protein | 36779 | 37714 | 936 | + |
| PCH45_039 | Putative RNA polymerase beta subunit | 37725 | 39332 | 1608 | + |
| PCH45_040 | Hypothetical protein | 39363 | 41426 | 2064 | + |
| PCH45_041 | Structural protein | 44314 | 41486 | 2829 | - |
| PCH45_042 | Structural protein | 44339 | 46684 | 2346 | + |
| PCH45_043 | Structural protein | 46697 | 47566 | 870 | + |
| PCH45_044 | Hypothetical protein | 47610 | 48164 | 555 | + |
| PCH45_045 | Structural protein | 48259 | 49242 | 984 | + |
| PCH45_046 | Hypothetical protein | 49255 | 50172 | 918 | + |
| PCH45_047 | Hypothetical protein | 50250 | 50606 | 357 | + |
| PCH45_048 | Hypothetical protein | 50619 | 52094 | 1476 | + |
| PCH45_049 | Crossover junction endodeoxyribonuclease RuvC | 52174 | 52773 | 600 | + |
| PCH45_050 | Thymidylate kinase | 52789 | 53466 | 678 | + |
| PCH45_051 | Hypothetical protein | 53447 | 53950 | 504 | + |
| PCH45_052 | Hypothetical protein | 53999 | 54742 | 744 | + |
| PCH45_053 | Hypothetical protein | 54866 | 55135 | 270 | + |
| PCH45_054 | Hypothetical protein | 55721 | 55185 | 537 | - |
| PCH45_055 | Structural protein | 56983 | 55718 | 1266 | - |
| PCH45_056 | Structural protein | 57044 | 59512 | 2469 | + |

| PCH45 ORF name | Protein function | Start (bp) | Stop (bp) | Length (bp) | Strand |
|---|---|---|---|---|---|
| PCH45_057 | Phage tail assembly | 59578 | 60057 | 480 | + |
| PCH45_058 | Structural protein | 60067 | 62730 | 2664 | + |
| PCH45_059 | Structural protein | 62747 | 63808 | 1062 | + |
| PCH45_060 | Long tail fiber, proximal subunit | 63821 | 65551 | 1731 | + |
| PCH45_061 | Hypothetical protein | 65596 | 66708 | 1113 | + |
| PCH45_062 | Structural protein | 67749 | 66757 | 993 | - |
| PCH45_063 | DNA double-strand break repair Rad50 ATPase | 67819 | 70281 | 2463 | + |
| PCH45_064 | Hypothetical protein | 70971 | 70321 | 651 | - |
| PCH45_065 | Phage protein CDS | 71600 | 70989 | 612 | - |
| PCH45_066 | Hypothetical protein | 72264 | 71611 | 654 | - |
| PCH45_067 | Structural protein | 73579 | 72266 | 1314 | - |
| PCH45_068 | Hypothetical protein | 73634 | 74158 | 525 | + |
| PCH45_069 | Ribonuclease H | 74210 | 75778 | 1569 | + |
| PCH45_070 | Hypothetical protein | 75775 | 76134 | 360 | + |
| PCH45_071 | Hypothetical protein | 77033 | 76179 | 855 | - |
| PCH45_072 | UvsX | 77114 | 78550 | 1437 | + |
| PCH45_073 | Phage protein | 78565 | 78852 | 288 | + |
| PCH45_074 | Structural protein | 79233 | 79910 | 678 | + |
| PCH45_075 | Hypothetical protein | 79913 | 80470 | 558 | + |
| PCH45_076 | Hypothetical protein | 80463 | 81263 | 801 | + |
| PCH45_077 | Hypothetical protein | 81784 | 81260 | 525 | - |
| PCH45_078 | Hypothetical protein | 81863 | 82384 | 522 | + |
| PCH45_079 | Hypothetical protein | 82548 | 83165 | 618 | + |
| PCH45_080 | Hypothetical protein | 83152 | 84246 | 1095 | + |
| PCH45_081 | Structural protein | 85080 | 84304 | 777 | - |
| PCH45_082 | Hypothetical protein | 85804 | 85088 | 717 | - |
| PCH45_083 | Hypothetical protein | 86531 | 85806 | 726 | - |
| PCH45_084 | Hypothetical protein | 88031 | 86544 | 1488 | - |
| PCH45_085 | Hypothetical protein | 88254 | 88021 | 234 | - |

| **PCH45 ORF name** | **Protein function** | **Start (bp)** | **Stop (bp)** | **Length (bp)** | **Strand** |
|---|---|---|---|---|---|
| PCH45_086 | RNA polymerase beta subunit | 92918 | 88308 | 4611 | - |
| PCH45_087 | Putative DNA-directed RNA polymerase beta subunit | 94552 | 92915 | 1638 | - |
| PCH45_088 | Phage tail fibers | 94578 | 101402 | 6825 | + |
| PCH45_089 | Structural protein | 101448 | 103802 | 2355 | + |
| PCH45_090 | Hypothetical protein | 103879 | 104571 | 693 | + |
| PCH45_091 | O-mannosyltransferase Ogm1 | 104581 | 105420 | 840 | + |
| PCH45_092 | Hypothetical protein | 105514 | 105954 | 441 | + |
| PCH45_093 | Polysaccharide deacetylase | 106039 | 106581 | 543 | + |
| PCH45_094 | Putative AraC-type DNA-binding protein | 106644 | 107222 | 579 | + |
| PCH45_095 | Hypothetical protein | 107233 | 107643 | 411 | + |
| PCH45_096 | Hypothetical protein | 107765 | 108397 | 633 | + |
| PCH45_097 | Hypothetical protein | 108402 | 108968 | 567 | + |
| PCH45_098 | Hypothetical protein | 109054 | 109653 | 600 | + |
| PCH45_099 | Hypothetical protein | 109771 | 110103 | 333 | + |
| PCH45_100 | Hypothetical protein | 110120 | 111019 | 900 | + |
| PCH45_101 | Hypothetical protein | 111024 | 111173 | 150 | + |
| PCH45_102 | Galactose oxidase | 111272 | 112207 | 936 | + |
| PCH45_103 | Hypothetical protein | 112276 | 112629 | 354 | + |
| PCH45_104 | Hypothetical protein | 112629 | 113093 | 465 | + |
| PCH45_105 | Putative phosphoesterase | 113093 | 114067 | 975 | + |
| PCH45_106 | Hypothetical protein | 114342 | 115454 | 1113 | + |
| PCH45_107 | Tail assembly-like protein | 115898 | 115500 | 399 | - |
| PCH45_108 | Putative lectin | 116633 | 115932 | 702 | - |
| PCH45_109 | Structural protein | 117332 | 116643 | 690 | - |
| PCH45_110 | Phage protein | 118061 | 117396 | 666 | - |
| PCH45_111 | Phage tail fiber protein | 121585 | 118115 | 3471 | - |
| PCH45_112 | Hypothetical protein | 121668 | 122072 | 405 | + |
| PCH45_113 | dUTP diphosphatase | 122139 | 122933 | 795 | + |
| PCH45_114 | Hypothetical protein | 122936 | 123214 | 279 | + |

| PCH45 ORF name | Protein function | Start (bp) | Stop (bp) | Length (bp) | Strand |
|---|---|---|---|---|---|
| PCH45_115 | Addiction module antidote protein, HigA family | 123263 | 123592 | 330 | + |
| PCH45_116 | Hypothetical protein | 123579 | 124001 | 423 | + |
| PCH45_117 | Hypothetical protein | 124011 | 124262 | 252 | + |
| PCH45_118 | Hypothetical protein | 124259 | 124696 | 438 | + |
| PCH45_119 | Ribonucleotide reductase of class Ia (aerobic), beta subunit | 124783 | 125889 | 1107 | + |
| PCH45_120 | Hypothetical protein | 125899 | 126246 | 348 | + |
| PCH45_121 | Hypothetical protein | 127112 | 126336 | 777 | - |
| PCH45_122 | Hypothetical protein | 127302 | 127892 | 591 | + |
| PCH45_123 | Hypothetical protein | 127902 | 128264 | 363 | + |
| PCH45_124 | Hypothetical protein | 128271 | 128615 | 345 | + |
| PCH45_125 | Hypothetical protein | 129006 | 129410 | 405 | + |
| PCH45_126 | Hypothetical protein | 129410 | 130141 | 732 | + |
| PCH45_127 | Phage protein | 130141 | 130764 | 624 | + |
| PCH45_128 | Hypothetical protein | 130782 | 131156 | 375 | + |
| PCH45_129 | Hypothetical protein | 131156 | 131542 | 387 | + |
| PCH45_130 | DNA recombination-mediator protein A | 131551 | 132114 | 564 | + |
| PCH45_131 | Hypothetical protein | 132125 | 133153 | 1029 | + |
| PCH45_132 | Hypothetical protein | 133153 | 133551 | 399 | + |
| PCH45_133 | Thymidylate synthase | 133725 | 134897 | 1173 | + |
| PCH45_134 | Putative Integrase | 134960 | 135301 | 342 | + |
| PCH45_135 | Phage protein | 135353 | 137023 | 1671 | + |
| PCH45_136 | Hypothetical protein | 137026 | 137364 | 339 | + |
| PCH45_137 | Hypothetical protein | 137381 | 137782 | 402 | + |
| PCH45_138 | Hypothetical protein | 137779 | 138108 | 330 | + |
| PCH45_139 | Metallophosphoesterase | 138690 | 138154 | 537 | - |
| PCH45_140 | Structural protein | 138743 | 139594 | 852 | + |
| PCH45_141 | Phage tail fibers | 139613 | 143611 | 3999 | + |
| PCH45_142 | Hypothetical protein | 143745 | 144353 | 609 | + |
| PCH45_143 | Putative endonuclease | 144355 | 144927 | 573 | + |

| PCH45 ORF name | Protein function | Start (bp) | Stop (bp) | Length (bp) | Strand |
|---|---|---|---|---|---|
| PCH45_144 | Hypothetical protein | 145018 | 145458 | 441 | + |
| PCH45_145 | Hypothetical protein | 145458 | 145964 | 507 | + |
| PCH45_146 | Hypothetical protein | 146249 | 146683 | 435 | + |
| PCH45_147 | Hypothetical protein | 146693 | 146968 | 276 | + |
| PCH45_148 | Hypothetical protein | 147016 | 147306 | 291 | + |
| PCH45_149 | Phage protein | 147303 | 147845 | 543 | + |
| PCH45_150 | Hypothetical protein | 147820 | 148422 | 603 | + |
| PCH45_151 | Hypothetical protein | 148434 | 148811 | 378 | + |
| PCH45_152 | Putative lytic transglycosylase | 148875 | 149624 | 750 | + |
| PCH45_153 | Hypothetical protein | 149705 | 150385 | 681 | + |
| PCH45_154 | Putative major structural protein | 151304 | 150438 | 867 | - |
| PCH45_155 | Putative tail sheath | 153375 | 151297 | 2079 | - |
| PCH45_156 | Putative structural protein | 153480 | 154409 | 930 | + |
| PCH45_157 | Putative structural protein | 154420 | 156954 | 2535 | + |
| PCH45_158 | Putative structural protein | 156957 | 158687 | 1731 | + |
| PCH45_159 | Phage protein | 158737 | 160845 | 2109 | + |
| PCH45_160 | Putative phosphohydrolase | 160929 | 161675 | 747 | + |
| PCH45_161 | Hypothetical protein | 161662 | 162174 | 513 | + |
| PCH45_162 | Phage protein | 163600 | 162377 | 1224 | - |
| PCH45_163 | Hypothetical protein | 163678 | 164328 | 651 | + |
| PCH45_164 | Hypothetical protein | 164472 | 164687 | 216 | + |
| PCH45_165 | Hypothetical protein | 164815 | 165219 | 405 | + |
| PCH45_166 | Hypothetical protein | 165432 | 165629 | 198 | + |
| PCH45_167 | Hypothetical protein | 165613 | 165840 | 228 | + |
| PCH45_168 | Hypothetical protein | 165905 | 167047 | 1143 | + |
| PCH45_169 | Hypothetical protein | 167059 | 167271 | 213 | + |
| PCH45_170 | Hypothetical protein | 167256 | 167534 | 279 | + |
| PCH45_171 | Hypothetical protein | 167633 | 167932 | 300 | + |
| PCH45_172 | Hypothetical protein | 168150 | 168935 | 786 | + |

| PCH45 ORF name | Protein function | Start (bp) | Stop (bp) | Length (bp) | Strand |
|---|---|---|---|---|---|
| PCH45_173 | Phage protein | 168960 | 169412 | 453 | + |
| PCH45_174 | Hypothetical protein | 169493 | 170200 | 708 | + |
| PCH45_175 | Hypothetical protein | 170330 | 170827 | 498 | + |
| PCH45_176 | Hypothetical protein | 170862 | 171200 | 339 | + |
| PCH45_177 | Hypothetical protein | 171202 | 171567 | 366 | + |
| PCH45_178 | Hypothetical protein | 171671 | 172390 | 720 | + |
| PCH45_179 | ATP synthase subunit I | 172391 | 172876 | 486 | + |
| PCH45_180 | Tail fiber protein | 172943 | 173602 | 660 | + |
| PCH45_181 | Hypothetical protein | 173658 | 174155 | 498 | + |
| PCH45_182 | Hypothetical protein | 174170 | 174853 | 684 | + |
| PCH45_183 | Phage protein | 174913 | 175485 | 573 | + |
| PCH45_184 | Hypothetical protein | 175518 | 176054 | 537 | + |
| PCH45_185 | Hypothetical protein | 176059 | 176499 | 441 | + |
| PCH45_186 | Phage protein | 177383 | 176547 | 837 | - |
| PCH45_187 | Putative tubulin-like protein | 177484 | 178458 | 975 | + |
| PCH45_188 | Hypothetical protein | 178656 | 178931 | 276 | + |
| PCH45_189 | Hypothetical protein | 179403 | 179870 | 468 | + |
| PCH45_190 | Hypothetical protein | 180107 | 180616 | 510 | + |
| PCH45_191 | Hypothetical protein | 180594 | 181085 | 492 | + |
| PCH45_192 | Hypothetical protein | 181165 | 181431 | 267 | + |
| PCH45_193 | Hypothetical protein | 181434 | 181817 | 384 | + |
| PCH45_194 | Hypothetical protein | 181814 | 182167 | 354 | + |
| PCH45_195 | Hypothetical protein | 182154 | 182627 | 474 | + |
| PCH45_196 | Phage protein | 182629 | 183009 | 381 | + |
| PCH45_197 | Hypothetical protein | 183006 | 183320 | 315 | + |
| PCH45_198 | Hypothetical protein | 183331 | 183747 | 417 | + |
| PCH45_199 | Putative T4-like DNA polymerase | 183878 | 186079 | 2202 | + |
| PCH45_200 | Phage protein | 187124 | 186039 | 1086 | - |
| PCH45_201 | Hypothetical protein | 187228 | 187869 | 642 | + |

| PCH45 ORF name | Protein function | Start (bp) | Stop (bp) | Length (bp) | Strand |
|---|---|---|---|---|---|
| PCH45_202 | Shell protein | 188038 | 190017 | 1980 | + |
| PCH45_203 | DNA-directed RNA polymerase beta prime subunit | 190108 | 191532 | 1425 | + |
| PCH45_204 | Hypothetical protein | 191529 | 192146 | 618 | + |
| PCH45_205 | Hypothetical protein | 192214 | 192606 | 393 | + |
| PCH45_206 | Hypothetical protein | 192599 | 193273 | 675 | + |
| PCH45_207 | Putative nuclease SbcCD D subunit | 193254 | 194438 | 1185 | + |
| PCH45_208 | Phage protein | 194562 | 195392 | 831 | + |
| PCH45_209 | Hypothetical protein | 195454 | 196257 | 804 | + |
| PCH45_210 | Phage protein | 196508 | 198061 | 1554 | + |
| PCH45_211 | Phage protein | 198074 | 199543 | 1470 | + |
| PCH45_212 | Polyribonucleotide nucleotidyltransferase | 199565 | 200986 | 1422 | + |
| PCH45_213 | Hypothetical protein | 201022 | 201528 | 507 | + |
| PCH45_214 | Hypothetical protein | 201525 | 201938 | 414 | + |
| PCH45_215 | Phage protein | 202330 | 201986 | 345 | - |
| PCH45_216 | Putative DNA directed RNA polymerase beta subunit | 202392 | 204557 | 2166 | + |
| PCH45_217 | Putative ATP-dependent DNA helicase | 204558 | 206561 | 2004 | + |
| PCH45_218 | Putative helicase | 206624 | 208150 | 1527 | + |
| PCH45_219 | Hypothetical protein | 208228 | 208566 | 339 | + |
| PCH45_220 | Phage protein | 208566 | 209045 | 480 | + |
| PCH45_221 | Phage protein | 209042 | 209254 | 213 | + |
| PCH45_222 | Hypothetical protein | 209368 | 209994 | 627 | + |
| PCH45_223 | Hypothetical protein | 210407 | 210042 | 366 | - |
| PCH45_224 | Phage protein | 212242 | 210428 | 1815 | - |
| PCH45_225 | Hypothetical protein | 212776 | 212252 | 525 | - |

**Table S2.** Spacer-target hits matching nucleoid forming phages.

| System type | Representative host assembly | Representative host strain | Representative viral target | Spacer-protospacer score | Target length | Spacer sequence | Representative shell homologue | Representative tubulin homologue |
|---|---|---|---|---|---|---|---|---|
| Type I-E | GCF_002941105 | Pseudomonas oleovorans strain YKJ | Cluster_sg_4850 | 32 | 253747 | TTCGACAGTGCGCCACCGGTTTCCTCTTTGGC | 3300011868_____Ga0122173_100014_____Ga0122173_100014162 | 3300011868_____Ga0122173_100014_____Ga0122173_100014181 |
| Type I-E | GCF_900455635 | Pseudomonas oleovorans strain NCTC10860 | Cluster_sg_4850 | 30 | 253747 | GTCGGTTTGAACAGGCGGTTGAGTTCGCGGAA | 3300011868_____Ga0122173_100014_____Ga0122173_100014162 | 3300011868_____Ga0122173_100014_____Ga0122173_100014181 |
| Type I-E | GCF_000953455 | Pseudomonas pseudoalcaligenes Ppseudo_Pac | Cluster_vOTU_002978 | 30 | 250515 | TACAACAAACCGGTCGAGCAGCAACTGCTTAA | 3300014697_____Ga0121494_100003_____Ga0121494_100003239;3300013343_____Ga0122025_100008_____Ga0122025_100008235 | 3300014697_____Ga0121494_100003_____Ga0121494_100003221;3300013343_____Ga0122025_100008_____Ga0122025_100008218 |
| Type I-E | GCF_003387475 | Kutzneria buriramensis strain DSM 45791 Ga0104555_123 | Cluster_sg_274913 | 25 | 313958 | GTGCGGCCGTTGTTGCCGAACGAGTTGAAGTTG | 3300013939_____Ga0117791_100022_____Ga0117791_100022225 | 3300013939_____Ga0117791_100022_____Ga0117791_100022237 |
| Type I-E | GCF_900455635 | Pseudomonas oleovorans strain NCTC10860 | Cluster_sg_6475;Cluster_sg_4850 | 25 | 276554 | GTCGGGTTGGGTAACGTTCAGAACTATGCGGTC | 3300014063_____Ga0122014_10008_____Ga0122014_1000854;3300011868_____Ga0122173_100014_____Ga0122173_100014162 | 3300014063_____Ga0122014_10008_____Ga0122014_1000876;3300011868_____Ga0122173_100014_____Ga0122173_100014181 |
| Type I-F | GCF_000793785 | Pseudomonas aeruginosa strain AZPAE14937 AZPAE14937_contig_14 | GCA_000866825.1 | 33 | 211215 | ACGATACTCTTACCGTCGGTACCGTCCTTACCG | CAG27117.1 | CAG27110.1 |
| Type I-F | GCF_003349755 | Vibrio cholerae strain OYP2A04 Vc_OYP2A04_Contig_21 | GCA_003441475.1 | 26 | 288967 | ACGATAGTACCAAGGTCACGCATGTTGCTGTT | AXH70812.1 | AXH70800.1 |
| Type III | GCF_003048445 | Campylobacter concisus strain P26UCO-S1 | Cluster_vOTU_000030 | 38 | 245889 | GATACTCCAGCATCAGCATTCATAGCAGTAGTGTTATA | 3300008515_____Ga0115189_1000003_____Ga0115189_100000365 | 3300008515_____Ga0115189_1000003_____Ga0115189_1000003113 |
| Type III | GCF_003048815 | Campylobacter concisus strain H9O-S1 | Cluster_vOTU_000030 | 38 | 243409 | AATCTTAGTAGGGATAGCTGGTTTCAAGAAGTTCTTAG | 3300008138_____Ga0114843_100237_____Ga0114843_1002374 | Inferred by other contigs in viral cluster |
| Type III | GCF_003048875 | Campylobacter concisus strain P11CDO-S1 | Cluster_vOTU_000030 | 37 | 243409 | AAAGATTTCTCTTTTACAGCTTCTTCTTTAGCAGCATTAGCTA | 3300006564_____Ga0100366_100027_____Ga0100366_10002761 | 3300006564_____Ga0100366_100027_____Ga0100366_10002712 |
| Type III | GCF_002912805 | Campylobacter concisus strain AAUH-10HCdes4 1103879 | Cluster_vOTU_000030 | 37 | 243409 | GTTATACATATAGCTTCTCTTATTGTAAACTGTAGCTTAGGAT | 3300007648_____Ga0105531_100012_____Ga0105531_100012149 | 3300007648_____Ga0105531_100012_____Ga0105531_100012100 |
| Type III | GCF_003048875 | Campylobacter concisus strain P11CDO-S1 | Cluster_vOTU_000030 | 37 | 243409 | GTAAGAGTATATACTGTAGATGAAACTAATGATATAC | 3300007648_____Ga0105531_100012_____Ga0105531_100012149 | 3300007648_____Ga0105531_100012_____Ga0105531_100012100 |
| Type III | GCF_006439235 | Haemophilus haemolyticus strain 60971 B Hi-3 Contig_2 | Cluster_vOTU_002372;Cluster_vOTU_019167 | 36 | 278993 | TCTAAGGTTGTCTTCTGATGTTCAGACATGCCCATC | 3300006545_____Ga0101078_100001_____Ga0101078_100001134;3300007931_____Ga0113985_100001_____Ga0113985_1000017;3300007932_____Ga0113995_100001_____Ga0113995_1000017 | 3300006545_____Ga0101078_100001_____Ga0101078_100001146;3300007931_____Ga0113985_100001_____Ga0113985_10000118;3300007932_____Ga0113995_100001_____Ga0113995_10000118 |
| Type III | GCF_003048445 | Campylobacter concisus strain P26UCO-S1 | Cluster_vOTU_000222 | 36 | 241426 | TCTACAGATGCATCATTAGGATCTGTTTTTAGACTAGC | 3300007220_____Ga0104054_100010_____Ga0104054_100010165 | 3300007220_____Ga0104054_100010_____Ga0104054_100010210 |
| Type III | GCF_900477945 | Haemophilus haemolyticus strain NCTC10839 | Cluster_vOTU_019167 | 35 | 278782 | TTACGGTAACCACGTTCAATAGTTTGGCTCATGTT | 3300007931_____Ga0113985_100001_____Ga0113985_1000017;3300007932_____Ga0113995_100001_____Ga0113995_1000017 | 3300007931_____Ga0113985_100001_____Ga0113985_10000118;3300007932_____Ga0113995_100001_____Ga0113995_10000118 |
| Type III | GCF_001949885 | Haemophilus parainfluenzae strain 65114 B Hi-3 Contig_3 | Cluster_vOTU_019167 | 35 | 278782 | TTGAGTTTGTTTAACTCTTTGATGATATCTTCGGC | 3300007931_____Ga0113985_100001_____Ga0113985_1000017;3300007932_____Ga0113995_100001_____Ga0113995_1000017 | 3300007931_____Ga0113985_100001_____Ga0113985_10000118;3300007932_____Ga0113995_100001_____Ga0113995_10000118 |
| Type III | GCF_003048445 | Campylobacter concisus strain P26UCO-S1 | Cluster_vOTU_000030 | 35 | 243409 | TATACTAGAAAGTTTACTAAAGTCGTGGTACTCTGGG | 3300006250_____Ga0099391_100017_____Ga0099391_10001759 | 3300006250_____Ga0099391_100017_____Ga0099391_100017105 |

| System type | Representative host assembly | Representative host strain | Representative viral target | Spacer-protospacer score | Target length | Spacer sequence | Representative shell homologue | Representative tubulin homologue |
|---|---|---|---|---|---|---|---|---|
| Type III | GCF_002912565 | Campylobacter concisus strain AAUH-10HCdes6 1106023 | Cluster_vOTU_000222 | 35 | 241426 | GATCTTTAATAAATTGTAACATGTTGAATCCTTTA | 3300007220_____Ga0104054_100010_____Ga0104054_100010165 | 3300007220_____Ga0104054_100010_____Ga0104054_100010210 |
| Type III | GCF_001008225 | Haemophilus haemolyticus strain 3P5 contig00093 | Cluster_vOTU_086883 | 35 | 220461 | TTGACATGTCGTACTCCTTTAAGGGTTAAAGCTTG | 3300007888_____Ga0111236_100001_____Ga0111236_100001143 | 3300007888_____Ga0111236_100001_____Ga0111236_100001162 |
| Type III | GCF_001949885 | Haemophilus parainfluenzae strain 65114 B Hi-3 Contig_3 | Cluster_vOTU_019167 | 34 | 278782 | AAGGTTTGGATCTTCTGGATCTTCAGAAGGGTTATT | 3300007931_____Ga0113985_100001_____Ga0113985_1000017;3300007932_____Ga0113995_100001_____Ga0113995_1000017 | 3300007931_____Ga0113985_100001_____Ga0113985_10000118;3300007932_____Ga0113995_100001_____Ga0113995_10000118 |
| Type III | GCF_002073715 | Neisseria sicca strain FDAARGOS_260 chromosome | Cluster_vOTU_003185 | 34 | 253014 | ACGAGTTTGTTGATTAATAGGCATTTTGATTTCC | 3300008130_____Ga0114850_100011_____Ga0114850_100011174 | 3300008130_____Ga0114850_100011_____Ga0114850_100011162 |
| Type III | GCF_003048445 | Campylobacter concisus strain P26UCO-S1 | Cluster_vOTU_000030 | 34 | 243409 | ATAAACTCATCTTCATCTTTTGGAAAATACTTATCAGCGA | 3300008136_____Ga0113979_100019_____Ga0113979_10001963 | 3300008136_____Ga0113979_100019_____Ga0113979_100019111 |
| Type III | GCF_900477945 | Haemophilus haemolyticus strain NCTC10839 | Cluster_vOTU_054382 | 34 | 241426 | TAACCGTGACGGGTAATCTTTTCAGATGAACCCG | 2124908010_____kg300_1843065_____kg300_01240230 | 2124908010_____kg300_1843065_____kg300_01240420 |
| Type III | GCF_001064745 | Morococcus cerebrosus strain 378_NMEN 905_1995_20947 | Cluster_vOTU_000365 | 34 | 241426 | CCATTCTGCAGTAGCTAATACTTCTTCTTGTTTA | 3300006743_____Ga0101795_100003_____Ga0101795_100003140;3300007980_____Ga0114366_1000001_____Ga0114366_100000176 | 3300006743_____Ga0101795_100003_____Ga0101795_100003128;3300007980_____Ga0114366_1000001_____Ga0114366_100000165 |
| Type III | GCF_002912805 | Campylobacter concisus strain AAUH-10HCdes4 1103879 | Cluster_vOTU_000222 | 34 | 241426 | GATCTTTAATAAATTGTAACATGTTGAATCCTTTAT | 3300007220_____Ga0104054_100010_____Ga0104054_100010165 | 3300007220_____Ga0104054_100010_____Ga0104054_100010210 |
| Type III | GCF_003044725 | Neisseria elongata strain C2013018262 C2013018262_S38_ctg_1733 | Cluster_sg_287804 | 34 | 241426 | TAGGATACCTGTACCGGATTTGAAACCAAACTTC | 3300008412_____Ga0115193_1000009_____Ga0115193_100000943 | 3300008412_____Ga0115193_1000009_____Ga0115193_100000932 |
| Type III | GCF_002990315 | Haemophilus influenzae strain 39P18H1 N39P18H1_8_2 | Cluster_vOTU_086883 | 34 | 220461 | TGATAAATGCCCTTCTCGCTGCACCATGTCTTTG | 3300007888_____Ga0111236_100001_____Ga0111236_100001143 | 3300007888_____Ga0111236_100001_____Ga0111236_100001162 |
| Type III | GCF_000222045 | Haemophilus haemolyticus M21127 M21127_015 | Cluster_vOTU_086883 | 34 | 220461 | AAGTCTTCTGGGGTAAAGCATTTGGCAATAATAA | 3300007888_____Ga0111236_100001_____Ga0111236_100001143 | 3300007888_____Ga0111236_100001_____Ga0111236_100001162 |
| Type III | GCF_000826045 | Haemophilus parahaemolyticus G321 CDBC01000003 | Cluster_vOTU_007652 | 34 | 211077 | TCAAACGGAGGAGGAGGGCTTCCTCCTCCTTTTT | 3300008643_____Ga0111423_100002_____Ga0111423_10000238 | 3300008643_____Ga0111423_100002_____Ga0111423_10000223 |
| Type III | GCF_003252925 | Haemophilus parahaemolyticus strain C2006000788 | Cluster_vOTU_007652 | 34 | 211077 | AACACGTGCCTGAAACGCTTCTAGAGCCTTTTGCAT | 3300008643_____Ga0111423_100002_____Ga0111423_10000238 | 3300008643_____Ga0111423_100002_____Ga0111423_10000223 |
| Type III | GCF_003494635 | Haemophilus haemolyticus strain M11818 | Cluster_vOTU_019167 | 33 | 278782 | TTAACTGTTTGACTTTATCCACCAATGGAGAGAAC | 3300007931_____Ga0113985_100001_____Ga0113985_1000017;3300007932_____Ga0113995_100001_____Ga0113995_1000017 | 3300007931_____Ga0113985_100001_____Ga0113985_10000118;3300007932_____Ga0113995_100001_____Ga0113995_10000118 |
| Type III | GCF_000287615 | Haemophilus sputorum HK 2154 ctg1200006325158 | Cluster_vOTU_011118 | 33 | 254212 | TATATTCATAACCTTCGAATTGCTGGATTGTTGTC | 3300008436_____Ga0115418_100017_____Ga0115418_10001719 | 3300008436_____Ga0115418_100017_____Ga0115418_10001732 |
| Type III | GCF_900477945 | Haemophilus haemolyticus strain NCTC10839 | Cluster_vOTU_009403 | 33 | 241426 | ATAAGCTGAAGAAGTCATCTGTTACGAATACACGT | 3300007789_____Ga0105762_100002_____Ga0105762_10000252 | 3300007789_____Ga0105762_100002_____Ga0105762_10000271 |
| Type III | GCF_006439235 | Haemophilus haemolyticus strain 60971 B Hi-3 Contig_2 | Cluster_vOTU_009403 | 33 | 241426 | TCACTCGAATGCCGAAATACGTCCGCAAGAAAT | 3300007789_____Ga0105762_100002_____Ga0105762_10000252 | 3300007789_____Ga0105762_100002_____Ga0105762_10000271 |
| Type III | GCF_001679135 | Haemophilus haemolyticus strain CCUG 24149 contig_22 | Cluster_vOTU_009403 | 33 | 241426 | CGTACTCCCTCTTCAGAGGTTTCTGTTGGTTTTGC | 3300007789_____Ga0105762_100002_____Ga0105762_10000252 | 3300007789_____Ga0105762_100002_____Ga0105762_10000271 |

1

| System type | Representative host assembly | Representative host strain | Representative viral target | Spacer-protospacer score | Target length | Spacer sequence | Representative shell homologue | Representative tubulin homologue |
|---|---|---|---|---|---|---|---|---|
| Type III | GCF_001679135 | Haemophilus haemolyticus strain CCUG 24149 contig_22 | Cluster_vOTU_009403 | 33 | 241426 | CACGAATCAATGGGTCACGTTGTGCATTAGTGT | 3300007789_____Ga0105762_100002_____Ga0105762_10000252 | 3300007789_____Ga0105762_100002_____Ga0105762_10000271 |
| Type III | GCF_001679135 | Haemophilus haemolyticus strain CCUG 24149 contig_22 | Cluster_vOTU_009403 | 33 | 241426 | ATTATCGTATATTTCACCTTTCTTTAGAGTGTATC | 3300007789_____Ga0105762_100002_____Ga0105762_10000252 | 3300007789_____Ga0105762_100002_____Ga0105762_10000271 |
| Type III | GCF_003048815 | Campylobacter concisus strain H9O-S1 | Cluster_vOTU_000030 | 33 | 239330 | TTGTTACCATAGACAACGAAGAGTGCTCTTTCATT | 3300008636_____Ga0111420_100018_____Ga0111420_100018146 | 3300008636_____Ga0111420_100018_____Ga0111420_10001899 |
| Type III | GCF_003252655 | Haemophilus haemolyticus strain C2001002324 C2001002324_S2_ctg_953 | Cluster_vOTU_086883 | 33 | 220461 | CACTGGTGCACGGTTGGTTTCAGACACTTCTACGT | 3300007888_____Ga0111236_100001_____Ga0111236_100001143 | 3300007888_____Ga0111236_100001_____Ga0111236_100001162 |
| Type III | GCF_001679135 | Haemophilus haemolyticus strain CCUG 24149 contig_22 | Cluster_vOTU_086883 | 33 | 220461 | AATAAATCGTATTCTTTACCTACTACTAATGGTTG | 3300007888_____Ga0111236_100001_____Ga0111236_100001143 | 3300007888_____Ga0111236_100001_____Ga0111236_100001162 |
| Type III | GCF_001679135 | Haemophilus haemolyticus strain CCUG 24149 contig_22 | Cluster_vOTU_086883 | 33 | 220461 | TCAACGGTATCAATACGATACAAAATAAATCTTTA | 3300007888_____Ga0111236_100001_____Ga0111236_100001143 | 3300007888_____Ga0111236_100001_____Ga0111236_100001162 |
| Type III | GCF_003048875 | Campylobacter concisus strain P11CDO-S1 | Cluster_vOTU_000030 | 32 | 243409 | GCTGGGGAATACTTATCGGCGAGATCCTTTATCTCATCTT | 3300006564_____Ga0100366_100027_____Ga0100366_10002761 | 3300006564_____Ga0100366_100027_____Ga0100366_10002712 |
| Type III | GCF_003048875 | Campylobacter concisus strain P11CDO-S1 | Cluster_vOTU_000030 | 32 | 243409 | TATACTTAATAGCTTTTCTCTGTCTGGTTTTCTAGAGA | 3300014024_____Ga0119801_100032_____Ga0119801_100032135 | Inferred by other contigs in viral cluster |
| Type III | GCF_002985245 | Haemophilus influenzae strain 39P1H1 N39P1H1_12_1 | Cluster_vOTU_009403 | 32 | 241426 | GATTTACGACCACCTTTATTTTGAGTTTGACGTA | 3300007789_____Ga0105762_100002_____Ga0105762_10000252 | 3300007789_____Ga0105762_100002_____Ga0105762_10000271 |
| Type III | GCF_001648245 | Eikenella corrodens strain NML04-0072 Eikcor_contig000033 | Cluster_vOTU_000638;Cluster_vOTU_003860;Cluster_vOTU_000593;Cluster_vOTU_021001 | 31 | 265823 | ATTTCTCTTTGGCTTTCTCAACGAACAAGGCTT | 3300008144_____Ga0114284_1000006_____Ga0114284_100000636;3300006247_____Ga0099374_1000004_____Ga0099374_100000413;3300008140_____Ga0114165_1000003_____Ga0114165_100000319;3300008480_____Ga0115173_1000002_____Ga0115173_100000261 | 3300008144_____Ga0114284_1000006_____Ga0114284_100000623;3300006247_____Ga0099374_1000004_____Ga0099374_1000004119;3300008140_____Ga0114165_1000003_____Ga0114165_1000003208;3300008480_____Ga0115173_1000002_____Ga0115173_100000274 |
| Type III | GCF_003048445 | Campylobacter concisus strain P26UCO-S1 | Cluster_vOTU_000030 | 31 | 243409 | TCTGTCTATATCGATAGACCTATCGACATTTACACTA | Inferred by other contigs in viral cluster | Inferred by other contigs in viral cluster |
| Type III | GCF_002990315 | Haemophilus influenzae strain 39P18H1 N39P18H1_8_2 | Cluster_vOTU_009403 | 31 | 241426 | CGATTTACGACCACCTTTATTTTGAGTTTGACGTA | 3300007789_____Ga0105762_100002_____Ga0105762_10000252 | 3300007789_____Ga0105762_100002_____Ga0105762_10000271 |
| Type III | GCF_001679135 | Haemophilus haemolyticus strain CCUG 24149 contig_22 | Cluster_vOTU_009403 | 31 | 241426 | TACCCGTAGGCTTTTCCATTAAGCCTTGTTCTT | 3300007789_____Ga0105762_100002_____Ga0105762_10000252 | 3300007789_____Ga0105762_100002_____Ga0105762_10000271 |
| Type III | GCF_004802095 | Haemophilus influenzae strain 60295_BAL_Hi1 60295_BAL_Hi1_7 | Cluster_vOTU_086883 | 31 | 220461 | CTGCAATGAAGTCATCTTGTTGTGGTTTACTTA | 3300007888_____Ga0111236_100001_____Ga0111236_100001143 | 3300007888_____Ga0111236_100001_____Ga0111236_100001162 |
| Type III | GCF_000222045 | Haemophilus haemolyticus M21127 M21127_015 | Cluster_vOTU_002372;Cluster_vOTU_019167 | 30 | 278993 | ACGCTGTGCATTCTTCCTGTACTTAATCTTGAGA | 3300006545_____Ga0101078_100001_____Ga0101078_100001134;3300007931_____Ga0113985_100001_____Ga0113985_1000017;3300007932_____Ga0113995_100001_____Ga0113995_1000017 | 3300006545_____Ga0101078_100001_____Ga0101078_100001146;3300007931_____Ga0113985_100001_____Ga0113985_10000118;3300007932_____Ga0113995_100001_____Ga0113995_10000118 |
| Type III | GCF_002912805 | Campylobacter concisus strain AAUH-10HCdes4 1103879 | Cluster_vOTU_000030 | 30 | 243409 | CCATTATAGTTAGCATTAAGTTTCTTAATATCTAGACCAC | 3300006250_____Ga0099391_100017_____Ga0099391_10001759 | 3300006250_____Ga0099391_100017_____Ga0099391_100017105 |
| Type III | GCF_003048445 | Campylobacter concisus strain P26UCO-S1 | Cluster_vOTU_000030 | 30 | 243409 | ATGTCATCTCGTTTACCATATCTTTAGTCATAGG | 3300006564_____Ga0100366_100027_____Ga0100366_10002761 | 3300006564_____Ga0100366_100027_____Ga0100366_10002712 |

1

| System type | Representative host assembly | Representative host strain | Representative viral target | Spacer-protospacer score | Target length | Spacer sequence | Representative shell homologue | Representative tubulin homologue |
|---|---|---|---|---|---|---|---|---|
| Type III | GCF_900454435 | Neisseria mucosa strain NCTC 10774 | Cluster_vOTU_000365;Cluster_vOTU_000639 | 30 | 241426 | GTAGTTATATTGTTTGTAATCAGCCATTTTTAAG | 3300006743_____Ga0101795_100003_____Ga0101795_100003140;3300007126_____Ga0102717_100009_____Ga0102717_100009191;3300007980_____Ga0114366_1000001_____Ga0114366_100000176;3300008082_____Ga0105968_1000005_____Ga0105968_100000519 | 3300006743_____Ga0101795_100003_____Ga0101795_100003128;3300007126_____Ga0102717_100009_____Ga0102717_100009203;3300007980_____Ga0114366_1000001_____Ga0114366_100000165;3300008082_____Ga0105968_1000005_____Ga0105968_1000005202 |
| Type III | GCF_001064475 | Morococcus cerebrosus strain 313_NMEN 1746_3886_34987 | Cluster_vOTU_000639 | 30 | 241426 | TCATGATTCTTATTCTTATGGGTATACAAGTGTA | 3300007126_____Ga0102717_100009_____Ga0102717_100009191;3300008082_____Ga0105968_1000005_____Ga0105968_100000519 | 3300007126_____Ga0102717_100009_____Ga0102717_100009203;3300008082_____Ga0105968_1000005_____Ga0105968_1000005202 |
| Type III | GCF_006439235 | Haemophilus haemolyticus strain 60971 B Hi-3 Contig_2 | Cluster_vOTU_009403 | 30 | 241426 | GGTTGTTTTCGATTTTGTTGATTAAGTTTTTCATGT | 3300007789_____Ga0105762_100002_____Ga0105762_10000252 | 3300007789_____Ga0105762_100002_____Ga0105762_10000271 |
| Type III | GCF_001679135 | Haemophilus haemolyticus strain CCUG 24149 contig_22 | Cluster_vOTU_086883 | 30 | 220461 | GTATTCACTTTCTGCACTATGGAAATATTCTAATCG | 3300007888_____Ga0111236_100001_____Ga0111236_100001143 | 3300007888_____Ga0111236_100001_____Ga0111236_100001162 |
| Type III | GCF_003048875 | Campylobacter concisus strain P11CDO-S1 | Cluster_vOTU_000030 | 29 | 243409 | AACTATTCTCTTAACTATGTCTTCTGTTAACGTACTCAT | 3300007648_____Ga0105531_100012_____Ga0105531_100012149;3300008636_____Ga0111420_100018_____Ga0111420_100018146 | 3300007648_____Ga0105531_100012_____Ga0105531_100012100;3300008636_____Ga0111420_100018_____Ga0111420_10001899 |
| Type III | GCF_003048445 | Campylobacter concisus strain P26UCO-S1 | Cluster_vOTU_000030 | 29 | 243409 | GATTGTACAGCTGACTCTGCATAAGCTTTAGAGAAAT | Inferred by other contigs in viral cluster | Inferred by other contigs in viral cluster |
| Type III | GCF_000023605 | Pectobacterium carotovorum subsp. carotovorum PC1 | Cluster_sg_274424 | 29 | 241426 | ACGTTATCCGCATACAGATCGAACTCCATCATG | 2029527006_____ACOFG988_contig16615_____ACOFGB_1405960 | 2029527006_____ACOFG988_contig16615_____ACOFGB_1405850 |
| Type III | GCF_003044725 | Neisseria elongata strain C2013018262 C2013018262_S38_ctg_1733 | Cluster_sg_287804 | 29 | 241426 | ACAATGAAATCCGTCTGACTGAGGTAGAAGCTAAA | 3300008412_____Ga0115193_1000009_____Ga0115193_100000943 | 3300008412_____Ga0115193_1000009_____Ga0115193_100000932 |
| Type III | GCF_003044725 | Neisseria elongata strain C2013018262 C2013018262_S38_ctg_1733 | Cluster_vOTU_026810 | 29 | 239603 | CCATGACGATCAACTCATCGCCGTCACCCAGTTTT | 3300007320_____Ga0104940_1000009_____Ga0104940_100000970 | 3300007320_____Ga0104940_1000009_____Ga0104940_100000980 |
| Type III | GCF_003044725 | Neisseria elongata strain C2013018262 C2013018262_S38_ctg_1733 | Cluster_vOTU_026810 | 29 | 239603 | TTCACCTTGATGCCTTTGGCGAAGTCGTATTCGAC | 3300007320_____Ga0104940_1000009_____Ga0104940_100000970 | 3300007320_____Ga0104940_1000009_____Ga0104940_100000980 |
| Type III | GCF_900477945 | Haemophilus haemolyticus strain NCTC10839 | Cluster_vOTU_086883 | 29 | 220461 | ACGTTGAACAATTGGCTTGAATTCCTCATAAAACT | 3300007888_____Ga0111236_100001_____Ga0111236_100001143 | 3300007888_____Ga0111236_100001_____Ga0111236_100001162 |
| Type III | GCF_001679135 | Haemophilus haemolyticus strain CCUG 24149 contig_22 | Cluster_vOTU_086883 | 29 | 220461 | ACTGCATCAATATTGCCATTATCGCACATTCTTGC | 3300007888_____Ga0111236_100001_____Ga0111236_100001143 | 3300007888_____Ga0111236_100001_____Ga0111236_100001162 |
| Type III | GCF_001644685 | Methylomonas sp. DH-1 | Cluster_sg_4850 | 28 | 253747 | CCCGACAGCCGGCGCATGGTAGTCAGCGCCTGGAATCC | 3300011868_____Ga0122173_100014_____Ga0122173_100014162 | 3300011868_____Ga0122173_100014_____Ga0122173_100014181 |
| Type III | GCF_002165875 | Campylobacter concisus strain Lasto220.96 Contig002 | Cluster_vOTU_004940 | 28 | 241426 | GTGTCGTAAATTTTATCTAAAATACTACTTACCGTTAT | 3300006460_____Ga0100061_100004_____Ga0100061_100004179 | 3300006460_____Ga0100061_100004_____Ga0100061_100004137 |
| Type III | GCF_001064745 | Morococcus cerebrosus strain 378_NMEN 905_1995_20947 | Cluster_vOTU_000365 | 28 | 241426 | TTTAGAATATCTTCTCTTTAAAAGAAATCAAG | 3300006743_____Ga0101795_100003_____Ga0101795_100003140;3300007980_____Ga0114366_1000001_____Ga0114366_100000176 | 3300006743_____Ga0101795_100003_____Ga0101795_100003128;3300007980_____Ga0114366_1000001_____Ga0114366_100000165 |

| System type | Representative host assembly | Representative host strain | Representative viral target | Spacer-protospacer score | Target length | Spacer sequence | Representative shell homologue | Representative tubulin homologue |
|---|---|---|---|---|---|---|---|---|
| Type III | GCF_900637305 | Cardiobacterium hominis strain NCTC10426 | Cluster_vOTU_005933;Cluster_sg_285216;Cluster_sg_292261;Cluster_vOTU_026988 | 27 | 269940 | CGACAAAATAGATATCTTCTTCTTGGAAGTCAG | 3300006498_____Ga0100374_100008_____Ga0100374_100008227;3300007368_____Ga0104977_1000002_____Ga0104977_100000212;3300011974_____Ga0119787_1000006_____Ga0119787_100000620;3300011925_____Ga0119789_100001_____Ga0119789_100001172 | 3300006498_____Ga0100374_100008_____Ga0100374_100008248;3300007368_____Ga0104977_1000002_____Ga0104977_1000002107;3300011974_____Ga0119787_1000006_____Ga0119787_1000006189;3300011925_____Ga0119789_100001_____Ga0119789_100001153 |
| Type III | GCF_000287615 | Haemophilus sputorum HK 2154 ctg120006325158 | Cluster_vOTU_011118 | 27 | 254212 | TTTTATCTTGTACGTTTGTAGGAGCAATTAC | 3300008436_____Ga0115418_100017_____Ga0115418_10001719 | 3300008436_____Ga0115418_100017_____Ga0115418_10001732 |
| Type III | GCF_900477945 | Haemophilus haemolyticus strain NCTC10839 | Cluster_vOTU_009403 | 27 | 241426 | TTGTAAGCACATGTTAACGATTAATGGTCCAGT | 3300007789_____Ga0105762_100002_____Ga0105762_10000252 | 3300007789_____Ga0105762_100002_____Ga0105762_10000271 |
| Type III | GCF_003048875 | Campylobacter concisus strain P11CDO-S1 | Cluster_vOTU_000030 | 27 | 239330 | GCTTACTAGCAATAAGCTAGCTATTAATACTCTTTTCATTATC | 3300008636_____Ga0111420_100018_____Ga0111420_100018146 | 3300008636_____Ga0111420_100018_____Ga0111420_10001899 |
| Type III | GCF_900477945 | Haemophilus haemolyticus strain NCTC10839 | Cluster_vOTU_086883 | 27 | 220461 | TTCGAAGCATGTACCAATAGCTCTTTTGGTAAT | 3300007888_____Ga0111236_100001_____Ga0111236_100001143 | 3300007888_____Ga0111236_100001_____Ga0111236_100001162 |

**Table S3.** Plasmid list

| Name | Features | Description | Construction | Reference |
|---|---|---|---|---|
| **Figure2.** | | | | |
| pPF1123 | RP4/oriT, pBR322_ori, CmR, mCherry, lacI/T5 | Naive vector, for plasmid interference assay | | Jackson *et al.*, 2019[11] |
| pPF1125 | RP4/oriT, pBR322_ori, CmR, mCherry, lacI/T5 | Phage priming vector with I-E PPS (spacer 1 CGT PAM) | derived from pPF1123 | Jackson *et al.*, 2019[11] |
| pPF1126 | RP4/oriT, pBR322_ori, CmR, mCherry, lacI/T5 | Phage priming vector with I-F PPS (spacer 2 GT PAM) | derived from pPF1123 | Jackson *et al.*, 2019[11] |
| pPF1255 | RP4/oriT, pBR322_ori, CmR, mCherry, lacI/T5 | Phage priming vector with I-E PPS (spacer 1 CGT PAM) with major capsid gene insert | PF2231/PF2232 into pPF1125 (SpeI, KpnI) | This study |
| pPF1256 | RP4/oriT, pBR322_ori, CmR, mCherry, lacI/T5 | Phage priming vector with I-F PPS (spacer 2 GT PAM) with major capsid gene insert | PF2231/PF2232 into pPF1126 (SpeI, KpnI) | This study |
| pPF974 | RP4/oriT, pBR322_ori, KmR, lacI/T5 | Type I-E repeat-BsaI-repeat construct for artificial crRNA | PF1962/PF1963 into pMAT16 (EcoRI, SphI) | Jackson *et al.*, 2019[11] |
| pPF975 | RP4/oriT, pBR322_ori, KmR, lacI/T5 | Type I-F repeat-BsaI-repeat construct for artificial crRNA | PF1964/PF1965 into pMAT16 (EcoRI, SphI) | Jackson *et al.*, 2019[11] |
| pPF976 | RP4/oriT, pBR322_ori, KmR, lacI/T5 | Type III-A repeat-BsaI-repeat construct for artificial crRNA | PF1981/PF1982 into pMAT16 (EcoRI, SphI) | This study |
| pPF1459 | RP4/oriT, pBR322_ori, KmR, lacI/T5 | anti-PCH45 I-E spacer overexpression. I-E_PCH45_PS12 | PF2811/ PF2812 into pPF974 (BsaI) | This study |
| pPF1460 | RP4/oriT, pBR322_ori, KmR, lacI/T5 | anti-PCH45 I-E spacer overexpression. I-E_PCH45_PS9 | PF2813/PF2814 into pPF974 (BsaI) | This study |
| pPF1461 | RP4/oriT, pBR322_ori, KmR, lacI/T5 | anti-PCH45 I-F spacer overexpression. I-F_PCH45-PS11 | PF2815/PF2816 into pPF975 (BsaI) | This study |
| pPF1462 | RP4/oriT, pBR322_ori, KmR, lacI/T5 | anti-PCH45 I-F spacer overexpression. I-F_PCH45-PS2 | PF2817/PF2818 into pPF975 (BsaI) | This study |
| pPF1467 | RP4/oriT, pBR322_ori, KmR, lacI/T5 | anti-PCH45 III-A spacer overexpression. III-A_PCH45_PS4 | PF2789/PF2790 into pPF976 (BsaI) | This study |
| pPF1443 | RP4/oriT, pBR322_ori, CmR, mCherry, lacI/T5 | Targeted vector with PCH45 capsid gene insert (*gp033_PCH45*), for plasmid interference assay | derived from pPF1255 (SpeI, SphI, MB nuclease followed by ligation) | This study |
| **Figure 3.** | | | | |
| **Spacer overexpression vectors** | | | | |
| pPF1465 | RP4/oriT, pBR322_ori, KmR, lacI/T5 | anti-PCH45 III-A spacer overexpression. III-A_PCH45_PS1 | PF2783/PF2784 into pPF976 (BsaI) | This study |
| pPF1466 | RP4/oriT, pBR322_ori, KmR, lacI/T5 | anti-PCH45 III-A spacer overexpression. III-A_PCH45_PS3 | PF2787/PF2788 into pPF976 (BsaI) | This study |
| pPF1468 | RP4/oriT, pBR322_ori, KmR, lacI/T5 | anti-PCH45 III-A spacer overexpression. III-A_PCH45_PS2 | PF2785/PF2786 into pPF976 (BsaI) | This study |
| pPF1469 | RP4/oriT, pBR322_ori, KmR, lacI/T5 | anti-PCH45 III-A spacer overexpression. III-A_PCH45_PS5 | PF2791/PF2792 into pPF976 (BsaI) | This study |
| pPF1470 | RP4/oriT, pBR322_ori, KmR, lacI/T5 | anti-PCH45 III-A spacer overexpression. III-A_PCH45_PS6 | PF2793/PF2794 into pPF976 (BsaI) | This study |
| pPF1994 | RP4/oriT, pBR322_ori, KmR, lacI/T5 | anti-PCH45 III-A spacer overexpression. III-A_PCH45_PS7 | PF3899/PF3900 into pPF976 (BsaI) | This study |
| pPF1995 | RP4/oriT, pBR322_ori, KmR, lacI/T5 | anti-PCH45 III-A spacer overexpression. III-A_PCH45_PS8 | PF3901/PF3902 into pPF976 (BsaI) | This study |
| pPF1996 | RP4/oriT, pBR322_ori, KmR, lacI/T5 | anti-PCH45 III-A spacer overexpression. III-A_PCH45_PS9 | PF3903/PF3904 into pPF976 (BsaI) | This study |

| Name | Features | Description | Construction | Reference |
|---|---|---|---|---|
| **Type III-A mutant construction** | | | | |
| pPF1117 | R6K_ori, RK2/OriT, SacB, MCR, CmR | Suicide vector for chromosomal knockouts | | Jackson *et al.*, 2019[11] |
| pPF1929 | R6K_ori, RK2/OriT, SacB, CmR, KmR | *cas7* knock out construct | Gibson assembly into pPF1117 (SalI, SphI). (PF3750-PF3585/PF3752-PF3753/PF3754-PF3751) | This study |
| pPF1930 | R6K_ori, RK2/OriT, SacB, CmR | *cas7*$^{D34A}$ knock in construct (KmR) | Gibson assembly into pPF1117 (SalI, SphI). (PF3750-PF3585/PF3589-PF3590/PF3755-PF3751) | This study |
| pPF1931 | R6K_ori, RK2/OriT, SacB, CmR | *cas7* wt knock in construct | Gibson assembly into pPF1117 (SalI, SphI). (PF3750-PF3751) | This study |
| pPF1932 | R6K_ori, RK2/OriT, SacB, CmR, KmR | Type III-A accessory nuclease knock out construct (KmR) | Gibson assembly into pPF1117 (SalI, SphI). (PF3743-PF3745/PF3746-PF3747/PF3748-PF3744) | This study |
| pPF1933 | R6K_ori, RK2/OriT, SacB, CmR | Type III-A accessory nuclease knock out construct (markless) | Gibson assembly into pPF1117 (SalI, SphI). (PF3743-PF3745/PF3749-PF3744) | This study |
| pPF1934 | R6K_ori, RK2/OriT, SacB, CmR | type III-A accessory nuclease wt knock in construct | Gibson assembly into pPF1117 (SalI, SphI). (PF3743-PF3744) | This study |
| pPF1935 | R6K_ori, RK2/OriT, SacB, CmR | *cas10* wt knock in construct | Gibson assembly into pPF1117 (SalI, SphI). (PF3756-PF3757) | This study |
| pPF1936 | R6K_ori, RK2/OriT, SacB, CmR | *cas10*$^{H17A,\ N18A,\ D618A\ D619A}$ (HD and Palm domain mutant) knock in construct | Gibson assembly into pPF1117 (SalI, SphI). (PF3756-PF2167/PF2166-PF2126/PF2127-PF3757) | This study |
| pPF1937 | R6K_ori, RK2/OriT, SacB, CmR | *cas10*$^{H17A,\ N18A}$ (HD domain mutant) knock in construct | Gibson assembly into pPF1117 (SalI, SphI). (PF3756-PF2167/PF2166-PF3757) | This study |
| pPF1938 | R6K_ori, RK2/OriT, SacB, CmR | *cas10*$^{D618A\ D619A}$ (Palm domain mutant) knock in construct | Gibson assembly into pPF1117 (SalI, SphI). (PF3756-PF2126/PF2127-PF3757) | This study |
| pPF781 | p15A_origin, OriT, AraC, MCR, CmR | Naive vector, for type III-A plasmid interference assay | pBAD30 backbone | Patterson *et al.*, 2016[24] |
| pPF1043 | p15A_origin, OriT, AraC, MCR, CmR, type III-A protospacer | Targeted vector with type III-A protospacer, for type III-A plasmid interference assay | pPF781 derivative | Patterson *et al.*, 2016[24] |
| **Fig 4.** | | | | |
| **Fluorescently tagged CRISPR-Cas complexes** | | | | |
| pPF1951 | R6K_ori, RK2/OriT, SacB, CmR | *cas10*USR-mCherry2 + linker-*cas10* fusion knock in construct | Gibson assembly into pPF1117 (SalI, SphI). (PF3813-PF3814/PF3811-PF3812/PF3815-PF3816) | This study |
| pPF1953 | R6K_ori, RK2/OriT, SacB, CmR | *cas8e*USR-mCherry2 + linker-*cas8e* fusion knock in construct | Gibson assembly into pPF1117 (SalI, SphI). (PF3817-PF3818/PF3811-PF3812/PF3819-PF3820) | This study |
| pPF1955 | R6K_ori, RK2/OriT, SacB, CmR | *cas8f*USR-mCherry2 + linker-*cas8f* fusion knock in construct | Gibson assembly into pPF1117 (SalI, SphI). (PF3821-PF3822/PF3811-PF3812/PF3823-PF3824) | This study |
| pQE80L-oriT stuffer | pBR322_ori, RP4/oriT, lacI/T5, AmpR | Expression vector induced with IPTG | | Watson *et al.*, unpublished[14] |

| Name | Features | Description | Construction | Reference |
|---|---|---|---|---|
| pPF1956 | pBR322_ori, RP4/oriT, lacI/T5, AmpR | mEGFP-shell gene fusion (*gp199_PCH45*) under T5 promoter | Gibson assembly into pQE80L-oriT stuffer (SphI, KpnI). (PF3825-PF3812/PF3826-PF3827) | This study |
| pPF1813 | R6K_ori, lacI/T5, TcR | Suicide vector with T5 promoter | | Watson *et al.*, unpublished[14] |
| pPF2036 | R6K_ori, lacI/T5, TcR | suicide vector with mCherry2 under T5 promoter | Gibson assembly into pPF1813 (EcoRI, XmaI). PF4005/PF4007 using gblock PF3810 as a template | This study |
| pPF1473 | RP4/oriT, pBR322_origin, KmR, lacI/T5 | anti-JS26 III-A spacer overexpression. III-A_JS26_PS4 (Helicase) | PF2759/PF2760 into pPF976 (BsaI) | This study |
| pPF1485 | RP4/oriT, pBR322_origin, KmR, lacI/T5 | anti-JS26 I-E spacer overexpression. I-E_JS26_PS20 (Tail length tape measure protein) | PF2797/PF2798 into pPF974 (BsaI) | This study |
| pPF1489 | RP4/oriT, pBR322_origin, KmR, lacI/T5 | anti-JS26 I-F spacer overexpression. I-F_JS26_PS73 (Tail length tape measure protein) | PF2801/PF2802 into pPF975 (BsaI) | This study |

### **Table S4.** Primer list

| Name | Sequence | Description/ Binding site | Notes |
|---|---|---|---|
| **General cloning** | | | |
| **PF2231** | TTTTACTAGTAGACGTTCAACAACGTCATG | PCH45 major capsid gene fwd | SpeI |
| **PF2232** | TTTTGGTACCGAAGTTATATTCGCGCGGTG | PCH45 major capsid gene rev | KpnI |
| **PF3809** | TCGTCTTCACCTCGAGAAATCAAAGAGGAGAAATTAACTATGGTGAGCAAGGGCGAGGAGCTGTTCACCGGGGTGGTGCCCATCCTGGTCGAGCTGGACGGCGACGTAAACGGCCACAAGTTCAGCGTGTCCGGCGAGGGCGAGGGCGATGCCACCTACGGCAAGCTGACCCTGAAGTTCATCTGCACCACCGGCAAGCTGCCCGTGCCCTGGCCCACCCTCGTGACCACCCTGACCTACGGCGTGCAGTGCTTCAGCCGCTACCCCGACCACATGAAGCAGCACGACTTCTTCAAGTCCGCCATGCCCGAAGGCTACGTCCAGGAGCGCACCATCTTCTTCAAGGACGACGGCAACTACAAGACCCGCGCCGAGGTGAAGTTCGAGGGCGACACCCTGGTGAACCGCATCGAGCTGAAGGGCATCGACTTCAAGGAGGACGGCAACATCCTGGGGCACAAGCTGGAGTACAACTACAACAGCCACAACGTCTATATCATGGCCGACAAGCAGAAGAACGGCATCAAGGTGAACTTCAAGATCCGCCACAACATCGAGGACGGCAGCGTGCAGCTCGCCGACCACTACCAGCAGAACACCCCCATCGGCGACGGCCCCGTGCTGCTGCCCGACAACCACTACCTGAGCACCCAGTCCAAGCTGAGCAAAGACCCCAACGAGAAGCGCGATCACATGGTCCTGCTGGAGTTCGTGACCGCCGCCGGGATCACTCTCGGCATGGACGAGCTGTACAAGGGCGGTGGAGGCGGATCCCCTGTTGATAGATCCAGTAATGAC | gblock RBS + mEGFP (no STOP codon) + linker(Gly5x-Ser) | |
| **PF3810** | TCGTCTTCACCTCGAGAAATCAAAGAGGAGAAATTAACTATGGTGAGCAAGGGCGAGGAGGATAACATGGCCATCATCAAGGAGTTCATGCGCTTCAAGGTGCACATGGAGGGCTCCGTGAACGGCCACGAGTTCGAGATCGAGGGCGAGGGCGAGGGCCGCCCCTACGAGGGCACCCAGACCGCCAAGCTGAAGGTGACCAAGGGTGGCCCCCTGCCCTTCGCCTGGGACATCCTGTCCCCTCAGTTCATGTACGGCTCCAAGGCCTACGTGAAGCACCCCGCCGACATCCCCGACTACTTGAAGCTGTCCTTCCCCGAGGGCTTCAATTGGGAGCGCGTGATGAACTTCGAGGACGGCGGCGTGGTGACCGTGACCCAGGACTCCTCCCTGCAGGACGGCGAGTTCATCTACAAGGTGAAGCTGCGCGGCACCAACTTCCCCTCCGACGGCCCCGTAATGCAGTGTCGTACCATGGGCTGGGAGGCCTCCACTGAGCGGATGTACCCCGAGGACGGCGCCCTGAAGGGCGAGATCAAGCAGAGGCTGAAGCTGAAGGACGGCGGCCACTACGACGCTGAGGTCAAGACCACCTACAAGGCCAAGAAGCCCGTGCAGCTGCCCGGCGCCTACAACGTCGACATCAAGTTGGACATCCTTTCCCACAACGAGGACTACACCATCGTGGAACAGTACGAACGCGCCGAGGGCCGCCACTCCACCGGCGGCATGGACGAGCTGTACAAGGGCGGTGGAGGCGGATCCCCTGTTGATAGATCCAGTAATGAC | RBS + mCherry2 (no STOP codon) + linker(Gly5x-Ser) gblock | |
| **PF3811** | ATGGTGAGCAAGGGCGAGGA | RBS + mEGFP/mCherry2-linker (gblockPF3809/gblockPF3810) fwd | |
| **PF3812** | GGATCCGCCTCCACCGCCC | RBS + mEGFP/mCherry2-linker (gblockPF3809/gblockPF3810) rev | |
| **PF3813** | TAGCTTGGCTGCAGGTCGACCAATCGGCCCCATAAATCAC | USR *cas10* fwd | overlap pPF1117, SalI |
| **PF3814** | TCCTCGCCCTTGCTCACCATTGACATCTCCTTGTGCCAAAAGG | USR *cas10* rev | overlap PF3811 |
| **PF3815** | AGGGCGGTGGAGGCGGATCCATGAACTGGCTTGCCGCCTC | DSR *cas10* fwd | overlap PF3812 |
| **PF3816** | GGGCTTCCCGGTATGCATGCGTCATCCACAGGCTGTCGAG | DSR *cas10* rev | overlap pPF1117, SphI |

| Name | Sequence | Description/ Binding site | Notes |
|---|---|---|---|
| **PF3817** | TAGCTTGGCTGCAGGTCGACCATTGCAGGATATCGCCAATCAG | USR *cas8e* fwd | overlap with pPF1117, SalI |
| **PF3818** | TCCTCGCCCTTGCTCACCATGGATCTATCTCCTCAGTTACACCTCG | USR *cas8e* rev | overlap PF3811 |
| **PF3819** | AGGGCGGTGGAGGCGGATCCATGTTTTCATTGATTGAAGCGCCG | DSR *cas8e* fwd | overlap pPF1117, SalI site |
| **PF3820** | GGGCTTCCCGGTATGCATGCCCCGGTACGATGCCCCACGC | DSR *cas8e* rev | overlap pPF1117, SphI |
| **PF3821** | TAGCTTGGCTGCAGGTCGACTTGCAACCGAAGCCACTGATC | USR *cas8f* fwd | overlap pPF1117, SalI |
| **PF3822** | TCCTCGCCCTTGCTCACCATTGCGCCTCCTGTTGTTATTGC | USR *cas8f* rev | overlap PF3811 |
| **PF3823** | AGGGCGGTGGAGGCGGATCCATGAAAGAAAACACATTGACGCAT | DSR *cas8f* fwd | overlap with pPF1117, with SalI |
| **PF3824** | GGGCTTCCCGGTATGCATGCGCCAACCATTGCGCCAGTTG | DSR *cas8f* rev | overlap pPF1117, SphI |
| **PF3825** | TCATACTAGGATCCGCATGCAAAGAGGAGAAATTAACTATGGTGAGCAAGGG | RBS + mEGFP/mCherry2 (gblock PF3809/PF3810) fwd | overlap pQE80L-stuffer, SphI |
| **PF3826** | AGGGCGGTGGAGGCGGATCCATGTCATTTAATGATAAAGAAAAACAAACTCCTG | Shell gene (gp199_PCH45) fwd | overlap PF3812 |
| **PF3827** | GCAGGTCGACCCGGGGTACCTTACCAACGGCCACCACGAC | Shell gene (gp199_PCH45) rev | overlap pPF1117, KpnI |
| **PF3828** | CCGTTGATAGGCATTTCGGC | Screening primer for mCherry2-*cas10* (in USR-DSR) fwd | |
| **PF3829** | GCCGGTGACGGGAAATCCTG | Screening primer for mCherry2-*cas10* (in USR-DSR) rev | |
| **PF3830** | GAACGGGCGGTATAGGCGGC | Screening primer for mCherry2-*cas10* (outside USD-DSR) fwd | |
| **PF3831** | GTGCGACGGCGAGCGCCGCC | Screening primer for mCherry2-*cas10* (outside USD-DSR) rev | |
| **PF3832** | GTGTCCTCACGGACGAGGTG | Screening primer for mCherry2-*cas8e* (in USR-DSR) fwd | |
| **PF3833** | CACGCGGCCAGTCAAGATCA | Screening primer for mCherry2-*cas8e* (in USR-DSR) rev | |
| **PF3834** | CGGAAGATGCCCGCATGCTG | Screening primer for mCherry2-*cas8e* (outside USD-DSR) fwd | |
| **PF3835** | CAGTACGCCTGTAGCTCATC | Screening primer for mCherry2-*cas8e* (outside USD-DSR) rev | |
| **PF3836** | GAACTGAGTTGGGTCAGCGC | Screening primer for mCherry2-*cas8f* (in USR-DSR) fwd | |
| **PF3837** | ATCCAGCTTGGCCTGCCGTC | Screening primer for mCherry2-*cas8f* (in USR-DSR) rev | |
| **PF3838** | CCGTCATCTCAGCGCTGCAC | Screening primer for mCherry2-*cas8f* (outside USD-DSR) fwd | |
| **PF3839** | GGCTGACGAACGGTGAAGAC | Screening primer for mCherry2-*cas8f* (outside USD-DSR) rev | |
| **PF3840** | GCTCGCCGACCACTACCAGC | Screening primer for mEGFP-shell fwd | |
| **PF3841** | GTCGTCACGACGGTTACGGC | Screening primer for mEGFP (in shell gene) rev | |
| **PF3842** | CCCACAACGAGGACTACACC | Screening primer for mCherry2 fwd | |
| **PF4005** | CAATTTCACACAGAATTCAAAGAGGAGAAATTAACTATGGTGAGCAAGG | mCherry2 into pPF1813 for plasmid chromosomal integration fwd | overlap pPF1813, EcoRI |
| **PF4007** | TTATTTGATGCCTCCCCGGGTTGTACAGCTCGTCCATGCCG | mCherry2 into pPF1813 for plasmid chromosomal integration rev | overlap pPF1813, XmaI |

| Name | Sequence | System | Description/ Binding site | Notes |
|---|---|---|---|---|
| **Construction of anti-PCH45 plasmid-borne mini-arrays** | | | | |
| **PF2811** | ACCGCCTTTCTCCAGATGGTCGATCGCGCTGTCAAAG | Type I-E | anti-PCH45 spacer I-E_PCH45_PS12 (anti-capsid_gene) fwd | BsaI |
| **PF2812** | AACACTTTGACAGCGCGATCGACCATCTGGAGAAAGG | Type I-E | anti-PCH45 spacer I-E_PCH45_PS12 (anti-capsid_gene) rev | BsaI |
| **PF2813** | ACCGTTTCTGTAGAAGCTGCAACCATTCTGGGCCTGG | Type I-E | anti-PCH45 spacer I-E_PCH45-PS9 (anti-capsid protein) fwd | BsaI |
| **PF2814** | AACACCAGGCCCAGAATGGTTGCAGCTTCTACAGAAA | Type I-E | anti-PCH45 spacer I-E_PCH45-PS9 (anti-capsid protein) rev | BsaI |
| **PF2815** | GAAAATCACCGGCGGTCTTAACGTTCTGCCGCAGAAG | Type I-F | anti-PCH45 spacer I-F_PCH45-PS11 (anti-capsid protein) fwd | BsaI |
| **PF2816** | TGAACTTCTGCGGCAGAACGTTAAGACCGCCGGTGAT | Type I-F | anti-PCH45 spacer I-F_PCH45-PS11 (anti-capsid protein) rev | BsaI |
| **PF2817** | GAAAATTCTGGGCCTGGCTCACAATGGCGTCGGCAAG | Type I-F | anti-PCH45 spacer I-F_PCH45_PS2 (anti-capsid protein) fwd | BsaI |
| **PF2818** | TGAACTTGCCGACGCCATTGTGAGCCAGGCCCAGAAT | Type I-F | anti-PCH45 spacer I-F_PCH45_PS2 (anti-capsid protein) rev | BsaI |
| **PF2783** | AGACGTAGAGTTGGTCAGGTCATCGCCTTTCAACTTACG | Type III-A | anti-PCH45 spacer III-A_PCH45_PS1 (anti- DNA polymerase) fwd | BsaI |
| **PF2784** | AGGACGTAAGTTGAAAGGCGATGACCTGACCAACTCTAC | Type III-A | anti-PCH45 spacer III-A_PCH45_PS1 (anti- DNA polymerase) rev | BsaI |
| **PF2785** | AGACCTTACGGATTTCTACTGGGACGCCATCGTACATGG | Type III-A | anti-PCH45 spacer III-A_PCH45_PS2 (anti-helicase) fwd | BsaI |
| **PF2786** | AGGACCATGTACGATGGCGTCCCAGTAGAAATCCGTAAG | Type III-A | anti-PCH45 spacer III-A_PCH45_PS2 (anti-helicase) rev | BsaI |
| **PF2787** | AGACCGGATAGACACCATTAGGAACGGTATTGTCATGCG | Type III-A | anti-PCH45 spacer III-A_PCH45_PS3 (anti-RNA polymerase) fwd | BsaI |
| **PF2788** | AGGACGCATGACAATACCGTTCCTAATGGTGTCTATCCG | Type III-A | anti-PCH45 spacer III-A_PCH45_PS3 (anti-RNA polymerase) rev | BsaI |
| **PF2789** | AGACGGTTTGCGCCCGGATGGAACTTCACGGTCATGTCG | Type III-A | anti-PCH45 spacer III-A_PCH45_PS4 (anti-capsid protein) fwd | BsaI |
| **PF2790** | AGGACGACATGACCGTGAAGTTCCATCCGGGCGCAAACC | Type III-A | anti-PCH45 spacer III-A_PCH45_PS4 (anti-capsid protein) rev | BsaI |
| **PF2791** | AGACCCACGTACCGTTAACGATGATTTTACGACCACGGG | Type III-A | anti-PCH45 spacer III-A_PCH45_PS5 (anti-terminase) fwd | BsaI |
| **PF2792** | AGGACCCGTGGTCGTAAAATCATCGTTAACGGTACGTGG | Type III-A | anti-PCH45 spacer III-A_PCH45_PS5 (anti-terminase) rev | BsaI |
| **PF2793** | AGACGTTTTTGTTACCGGCTTCGTTGTTGTCGTAGAACG | Type III-A | anti-PCH45 spacer III-A_PCH45_PS6 (anti-TLP) fwd | BsaI |
| **PF2794** | AGGACGTTCTACGACAACAACGAAGCCGGTAACAAAAAC | Type III-A | anti-PCH45 spacer III-A_PCH45_PS6 (anti-TLP) rev | BsaI |
| **PF3899** | AGACGACATGACCGTGAAGTTCCATCCGGGCGCAAACCG | Type III-A | anti-PCH45 spacer III-A_PCH45_PS7 (reverse PS4_major capsid gene) fwd | BsaI |
| **PF3900** | AGGACGGTTTGCGCCCGGATGGAACTTCACGGTCATGTC | Type III-A | anti-PCH45 spacer III-A_PCH45_PS7 (reverse PS4_major capsid gene) rev | BsaI |
| **PF3901** | AGACCCGTGGTCGTAAAATCATCGTTAACGGTACGTGGG | Type III-A | anti-PCH45 spacer III-A_PCH45_PS8 (reverse PS5_terminase) fwd | BsaI |
| **PF3902** | AGGACCCACGTACCGTTAACGATGATTTTACGACCACGG | Type III-A | anti-PCH45 spacer III-A_PCH45_PS8 (reverse PS5_terminase) rev | BsaI |
| **PF3903** | AGACGTTCTACGACAACAACGAAGCCGGTAACAAAAACG | Type III-A | anti-PCH45 spacer III-A_PCH45_PS9 (reverse PS6_tubulin-like gene) fwd | BsaI |
| **PF3904** | AGGACGTTTTTGTTACCGGCTTCGTTGTTGTCGTAGAAC | Type III-A | anti-PCH45 spacer III-A_PCH45_PS9 (reverse PS6_tubulin-like gene) rev | BsaI |

| Name | Sequence | System | Description/ Binding site | Notes |
|---|---|---|---|---|
| **PF2759** | AGAC**ATACAGTTTTTCGTAATAGTCCTCGTTGGCCTGC**G | Type III-A | F cloning anti-JS26 spacer III-A_JS26_PS4 (anti-helicase) fwd | BsaI |
| **PF2760** | AGGAC**GCAGGCCAACGAGGACTATTACGAAAAACTGTAT** | Type III-A | R cloning anti-JS26 spacer III-A_JS26_PS4 (anti-helicase) rev | BsaI |
| **PF2797** | ACCG**GCCGCTTTGTTGGCCTCCGCAAACTGGCGAAT**G | Type I-E | F cloning anti-JS26 spacer I-E_JS26_PS20 (anti-TMP) fwd | BsaI |
| **PF2798** | AACAC**ATTCGCCAGTTTGCGGAGGCCAACAAAGCGGC** | Type I-E | R cloning anti-JS26 spacer I-E_JS26_PS20 (anti-TMP) rev | BsaI |
| **PF2801** | GAAA**GGTAATGGCACCCATGTCGCCCTGCAGCGCTT**G | Type I-F | F cloning anti-JS26 spacer I-F_JS26_PS73 (anti-TMP) fwd | BsaI |
| **PF2802** | TGAAC**AAGCGCTGCAGGGCGACATGGGTGCCATTACC** | Type I-F | R cloning anti-JS26 spacer I-F_JS26_PS73 (anti-TMP) rev | BsaI |
| **CRISPR expansion PCRs** | | | | |
| **PF1989** | TAAGTTAGTGTTCTTTAACAAGCAGGA | Type I-E | *Serratia* I-E CRISPR expansion screening fwd | |
| **PF1887** | GTTAAGTCAGCAGGCGTTTAGTCG | Type I-E | *Serratia* I-E CRISPR expansion screening rev | |
| **PF1990** | CACGAAAATGATAATTGATGCTGAT | Type I-F | *Serratia* I-F CRISPR expansion screening fwd | |
| **PF1888** | CATCTGATGCTGACGACACTG | Type I-F | *Serratia* I-F CRISPR expansion screening rev | |
| **Mutant construction** | | | | |
| **PF3589** | CATACGCCGCGACTGACAAG | Type III-A | *cas7*$^{D34A}$ fwd | |
| **PF3590** | CTCAAAGCGCGGCTGGAGATC | Type III-A | *cas7*$^{D34A}$ Rev | |
| **PF3591** | CATACGCCGCGACTGACAAGGACATCTCATGCAACTGAACAATATCCAGAC<br>GTTACGCGCCACTCTGGTGTGTGAAACCGGGTTACACATTGGCGGCGGCG<br>ACACCGCGTTGCAGATTGGCGGCATCGCTAGCGCCGTGGTGCGCCACCCG<br>TTGACCCAGCAACCTTACATTCCCGGCTCCAGCCTGAAAGGCAAACTGCGC<br>AGCCTGCTCGAATGGCGCGCTGGTGTGGTCGGCGATACCGAGGGCAAGGT<br>GCTCAGTCATCAGGTTTACCAGCAACTGATCGACGATAAAAAACAGGCACA<br>GGCACTACAGGCACTGAAAATCCTGCAACTGTTTGGCGTCAGCGGCGGCG<br>ACAAGCTTTCCGCCGAACAAGCCCAACAGATCGGCCCGACTCGCCTGTCCT<br>TCTGGGATTGCGAATTTGACGAACACTGGCTGGCGCAGCAAGGCGGGCGC<br>GTCCAGACCGAAGAGAAAGCGGAAAACTGTATTGACCGCATCAGTGGCGTG<br>GCGCTGCACCCGCGCTTTATCGAGCGTGTGCCCGCCGGCAGCCGCTTTGA<br>CTTTCGCCTCACCGTGCGCCAGCTTGATGGCGACAGCCCCGACCTGCTCG<br>ACACCCTGTTGGCGGGCCTGAAAATGCTGGAGCTGGACGGGCTAGGCGGC<br>AGCATTTCCCGTGGGTATGGCAAAGTGCGCTTTGAAGCGCTGACCCTCGAC<br>GGGAAAGATCTCCAGCCGCGCTTTGAGCAGTTACAGCCGTTTAAACACACC<br>ACGCAGGGAGCCGGATGACTGCAGCCTGTTGATAGATCCAGTAATGAC | Type III-A | *cas7*$^{D34A}$ gblock | |
| **PF3743** | TAGCTTGGCTGCAGGTCGACCAACTCTGTGGAGGAGGCCG | Type III-A | USR type III-A accessory nuclease fwd | overlap pPF1117, SalI |
| **PF3744** | GGGCTTCCCGGTATGCATGCCTTATGTGCTGGAGCCAGCG | Type III-A | DSR type III-A accessory nuclease rev | overlap pPF1117, SphI |
| **PF3745** | CAGCCCCCATTCAGGCACAG | Type III-A | USR type III-A accessory nuclease rev | |
| **PF3746** | CTGTGCCTGAATGGGGGCTGGGTGAGAATCCAGGGGTCCC | Type III-A | Kanamycin resistance cassette in pSEVA221 fwd (nuclease knock out vector) | overlap PF3745 |
| **PF3747** | TCCCCGCTAACGCCAGGAAGGGACAACGCGCGGACCG | Type III-A | Kanamycin resistance cassette in pSEVA221 rev (nuclease knock out vector) | overlap PF3748 |

| Name | Sequence | System | Description/ Binding site | Notes |
|---|---|---|---|---|
| **PF3748** | GTCCCTTCCTGGCGTTAGCGGGGA | Type III-A | DSR type III-A accessory nuclease fwd (knock out vector) | |
| **PF3749** | CTGTGCCTGAATGGGGGCTGCTTCCTGGCGTTAGCGGGGA | Type III-A | DSR type III-A accessory nuclease rev (knock out vector) | overlap PF3745 |
| **PF3750** | TAGCTTGGCTGCAGGTCGACCTCCTCCGGCCTGCTTTAC | Type III-A | USR *cas7* fwd | overlap pPF1117, SalI |
| **PF3585** | CTTGTCAGTCGCGGCGTATG | Type III-A | USR *cas7* rev | |
| **PF3751** | GGGCTTCCCGGTATGCATGCCTGTCGGCACTGTGCCAGTG | Type III-A | DRS *cas7* rev | overlap pPF1117, SphI |
| **PF3752** | CATACGCCGCGACTGACAAGGGTGAGAATCCAGGGGTCCC | Type III-A | Kanamycin resistance cassette in pSEVA221 fwd (*cas7* knock out vector) | overlap PF3585 |
| **PF3753** | CGGCTCCCTGCGTGGTGTGTGGACAACGCGCGGACCG | Type III-A | Kanamycin resistance cassette in pSEVA221 rev (*cas7* knock out vector) | overlap PF3754 |
| **PF3754** | TGTCCACACACCACGCAGGGAGC | Type III-A | DSR *cas7* fwd | |
| **PF3755** | GATCTCCAGCCGCGCTTTGAGACACACCACGCAGGGAGC | Type III-A | DSR *cas7* fwd | overlap PF3590 |
| **PF1934** | AGA**GTCGAC**CTTACACGGTTGCGCGTCTAAG | Type III-A | USR cas10 fwd | SalI |
| **PF1935** | CAC**GGATCC**ATGGCAAGAGGCGGCAAG | Type III-A | USR cas10 rev | BamHI |
| **PF1936** | TTT**GGATCC**GGCGAGTAAAACGCATAACAAC | Type III-A | DSR cas10 fwd | BamHI |
| **PF1937** | TTT**TCTAGAGCATGC**GTGTTCGTCAAATTCGCAATC | Type III-A | DSR cas10 rev | XbaI and SphI |
| **PF3756** | TAGCTTGGCTGCAGGTCGACCTTACACGGTTGCGCGTCTAAG | Type III-A | USR *cas10* fwd | overlap pPF1117, SalI |
| **PF3757** | GGGCTTCCCGGTATGCATGCGCAATCCCAGAAGGACAGGC | Type III-A | DSR *cas10* rev | overlap pPF1117, SphI |
| **PF2166** | TGCCTTTGCGCTGTTAGCTGCCCTGAAACCGCTGGCGCAGCGT | Type III-A | *cas10*$^{\text{H17A, N18A}}$ (HD domain mutant) fwd | overlap PF2167 |
| **PF2167** | AGCTAACAGCGCAAAGGCAGC | Type III-A | *cas10*$^{\text{H17A, N18A}}$ (HD domain mutant) rev | |
| **PF2126** | GGCGAACACGGTGTAGGTG | Type III-A | *cas10*$^{\text{D618A, D619A}}$ (Palm domain mutant) rev | |
| **PF2127** | CACCTACACCGTGTTCGCCGGCGGCGCAGCCTTCTTTTTGATTGGCCCGTG | Type III-A | *cas10*$^{\text{D618A, D619A}}$ (Palm domain mutant) fwd | overlap PF2126 |
| **PF3606** | GCTGTTATGACATGGGTTTG | Type III-A | Screening primer for hypothetical nuclease mutagenesis fwd | |
| **PF3607** | CATTGAGACCCAAGATAGCG | Type III-A | Screening primer for hypothetical nuclease mutagenesis rev | |
| **PF3608** | CATCTGCCTGTATAACGAGG | Type III-A | Screening primer for *cas7* mutagenesis fwd | |
| **PF3609** | CTGTCTGCCAGTAGCAGTGG | Type III-A | Screening primer for *cas7* mutagenesis rev | |
| **PF3610** | CAGTTTTCCGCTTTCTCTTC | Type III-A | Screening primer for *cas10* mutagenesis rev | |

**Table S5**. Bacterial strains used in this study

| Species | Strain | Description | Notes | Construction | Reference |
|---|---|---|---|---|---|
| *E. coli* | DH5α | Cloning strain | | | Gibco/BRL |
| *E. coli* | ST18 | Auxotrophic donor for biparental conjugation | Requires ALA | | Thoma and Schobert, 2009[25] |
| *Serratia* sp. ATCC 39006 | LacA | *lac* EMS mutant, denoted WT | | | Thomson *et al.*, 2000[26] |
| **Type III-A mutants** | | | | | |
| *Serratia* sp. ATCC 39006 | PCF303 | Δ*cas10* | Markerless | allelic exchange with pPF927 | This study |
| *Serratia* sp. ATCC 39006 | PCF682 | Δ*cas7* | KmR | allelic exchange with pPF1929 | This study |
| *Serratia* sp. ATCC 39006 | PCF683 | $cas7^{D34A}$ | | allelic exchange with pPF1930 in PF682 | This study |
| *Serratia* sp. ATCC 39006 | PCF684 | *cas7* wt | | allelic exchange with pPF1931 in PF682 | This study |
| *Serratia* sp. ATCC 39006 | PCF685 | Δ hypothetical nuclease | KmR | allelic exchange with pPF1932 | This study |
| *Serratia* sp. ATCC 39006 | PCF686 | Δ hypothetical nuclease | Markerless | allelic exchange with pPF1933 in PF685 | This study |
| *Serratia* sp. ATCC 39006 | PCF687 | hypothetical nuclease wt | | allelic exchange with pPF1934 in PCF303 | This study |
| *Serratia* sp. ATCC 39006 | PCF688 | *cas10* wt | | allelic exchange with pPF1935 in PCF303 | This study |
| *Serratia* sp. ATCC 39006 | PCF690 | $cas10^{H17A,\ N18A}$ (HD mutant) | | allelic exchange with pPF1937 in PCF303 | This study |
| *Serratia* sp. ATCC 39006 | PCF691 | $cas10^{D618A,\ D619A}$ (Palm mutant) | | allelic exchange with pPF1938 in PCF303 | This study |
| **Strains with spacers in native CRISPR arrays** | | | | | |
| *Serratia* sp. ATCC 39006 | PCF543 | anti-PCH45 spacer in type I-E system | anti-major capsid gene | plasmid loss with pPF1255 | This study |
| *Serratia* sp. ATCC 39006 | PCF544 | anti-PCH45 spacer in type I-E system | anti-major capsid gene | plasmid loss with pPF1255 | This study |
| *Serratia* sp. ATCC 39006 | PCF545 | anti-PCH45 spacer in type I-E system | anti-major capsid gene | plasmid loss with pPF1255 | This study |
| *Serratia* sp. ATCC 39006 | PCF547 | anti-PCH45 spacer in type I-F system | anti-major capsid gene | plasmid loss with pPF1256 | This study |
| *Serratia* sp. ATCC 39006 | PCF548 | anti-PCH45 spacer in type I-F system | anti-major capsid gene | plasmid loss with pPF1256 | This study |
| *Serratia* sp. ATCC 39006 | PCF591 | anti-PCH45 spacer in type I-E system | anti-major capsid gene | plasmid loss with pPF1255 | This study |
| *Serratia* sp. ATCC 39006 | PCF592 | anti-PCH45 spacer in type I-E system | anti-major capsid gene | plasmid loss with pPF1255 | This study |
| *Serratia* sp. ATCC 39006 | PCF593 | anti-PCH45 spacer in type I-E system | anti-major capsid gene | plasmid loss with pPF1255 | This study |
| **Fluorescently tagged interference complexes** | | | | | |
| *Serratia* sp. ATCC 39006 | PCF732 | Fluorescently tagged Type III-A interference complex (*mCherry2-cas10* fusion) | | Allelic exchange in lacA with pPF1951 | This study |

| | | | | | |
|---|---|---|---|---|---|
| *Serratia* sp. ATCC 39006 | PCF734 | Fluorescently tagged Type I-E interference complex (*mCherry2-cas8e* fusion) | | Allelic exchange in lacA with pPF1953 | This study |
| *Serratia* sp. ATCC 39006 | PCF736 | Fluorescently tagged Type I-F interference complex (*mCherry2-cas8f* fusion) | | Allelic exchange in lacA with pPF1955 | This study |
| *Serratia* sp. ATCC 39006 | PCF761 | Fluorescently tagged Type I-E interference complex (*mCherry2-cas8e*) under T5 promoter in chromosome | TcR, inducible with IPTG | pPF2038 chromosomal integration in PCF734 (upstream *cse* complex) | This study |
| *Serratia* sp. ATCC 39006 | PCF763 | Fluorescently tagged Type I-F interference complex (*mCherry2-cas8f*) under T5 promoter in chromosome | TcR, inducible with IPTG | pPF2040 chromosomal integration in PCF736 (upstream *csy* complex) | This study |
| *Serratia* sp. ATCC 39006 | PCF765 | Fluorescently tagged Type III-A interference complex (*mCherry2-cas10*) under T5 promoter in chromosome | TcR, inducible with IPTG | pPF2042 chromosomal integration in PCF732 (upstream *csm* complex) | This study |

**Table S6.** Spacer list in native CRISPR arrays

| Strain Name | System | No. of new spacers | Sequence | Protospacer | Strand |
|---|---|---|---|---|---|
| PCF544 | Type I-E | 5 | CTGTGAGCAGTGTTGAACTTCAGGTACTGAGT | Phage major capsid insert (*gp033_PCH45*) | - |
| | | | CTGTGACCGTCTCCGGGAGCTGCATGTGTCAG | plasmid backbone | - |
| | | | GCACAATTCTCATGTTTGACAGCTTATCATCG | plasmid backbone | - |
| | | | CAAATAAATTTTTTATGATTTCTCGAGCTCAT | plasmid backbone | - |
| | | | TCAGAGGTGGCGAAACCCGACAGGACTATAAA | pBR322origin | + |
| PCF545 | Type I-E | 4 | GGACTCCTCCTTTATTTTATTTAGAATTCTGT | plasmid backbone | - |
| | | | CGCATGAACTCCTTGATGATGGCCATGTTATC | *mCherry2* | - |
| | | | TTAGCTCACTCATTAGGCACAATTCTCATGTT | plasmid backbone | - |
| | | | TCCGTTTCCAGGATTTTGTGGTTCAGCAGCGC | Phage major capsid insert (*gp033_PCH45*) | - |
| PCF591 | Type I-E | 2 | TTTCTGTAGAAGCTGCAACCATTCTGGGCCTG | Phage major capsid insert (*gp033_PCH45*) | + |
| | | | CTGTCAAACATGAGAATTGTGCCTAATGAGTG | plasmid backbone | + |
| PCF592 | Type I-E | 1 | CCTTTCTCCAGATGGTCGATCGCGCTGTCAAA | Phage major capsid insert (*gp033_PCH45*) | - |
| PCF593 | Type I-E | 3 | GGTGCACCGAACTTCACCGACTCTCTGGACCG | Phage major capsid insert (*gp033_PCH45*) | + |
| | | | TTGGACATCACCTCCCACAACGAGGACTACAC | *mCherry2* | + |
| | | | CATTCTGCCGACATGGAAGCCATCACAAACGG | Undetermined target | |
| PCF547 | Type I-F | 1 | ATTCTGGGCCTGGCTCACAATGGCGTCGGCAA | Phage major capsid insert (*gp033_PCH45*) | + |
| PCF548 | Type I-F | 2 | GATAGCGGAACGGGAAGGCGACTGGAGTGCCA | *lacI* | - |
| | | | ATCACCGGCGGTCTTAACGTTCTGCCGCAGAA | Phage major capsid insert (*gp033_PCH45*) | + |

**Table S7.** Spacers expressed from mini-array in plasmid

| Spacers expressed from plasmids-borne mini arrays | | | | |
|---|---|---|---|---|
| **Plasmid Name** | **Spacer** | **System** | **Spacer Sequence** | **Target** |
| **pPF1459** | S1 | Type I-E | CCTTTCTCCAGATGGTCGATCGCGCTGTCAAA | Major capsid gene (*gp033*) |
| **pPF1460** | S4 | Type I-E | TTTCTGTAGAAGCTGCAACCATTCTGGGCCTG | Major capsid gene (*gp033*) |
| **pPF1461** | S8 | Type I-F | ATCACCGGCGGTCTTAACGTTCTGCCGCAGAA | Major capsid gene (*gp033*) |
| **pPF1462** | S2 | Type I-F | ATTCTGGGCCTGGCTCACAATGGCGTCGGCAA | Major capsid gene (*gp033*) |
| **pPF1468** | S12 | Type III-A | CTTACGGATTTCTACTGGGACGCCATCGTACATG | DNA helicase (*gp217*) |
| **pPF1466** | S9 | Type III-A | CGGATAGACACCATTAGGAACGGTATTGTCATGC | RNA polymerase beta subunit (*gp084*) |
| **pPF1467** | S3 | Type III-A | GGTTTGCGCCCGGATGGAACTTCACGGTCATGTC | Major capsid gene (*gp033*) |
| **pPF1469** | S10 | Type III-A | CCACGTACCGTTAACGATGATTTTACGACCACGG | Terminase large subunit (*gp159*) |
| **pPF1470** | S11 | Type III-A | GTTTTTGTTACCGGCTTCGTTGTTGTCGTAGAAC | Tubulin-like gene (*gp187*) |
| **pPF1473** | S15 | Type III-A | ATACAGTTTTTCGTAATAGTCCTCGTTGGCCTGC | DNA helicase (JS26) |
| **pPF1485** | S16 | Type I-E | GCCGCTTTGTTGGCCTCCGCAAACTGGCGAAT | Tail length tape measure gene (JS26) |
| **pPF1489** | S17 | Type I-F | GGTAATGGCACCCATGTCGCCCTGCAGCGCTT | Tail length tape measure gene (JS26) |
| **pPF1994** | S13 | Type III-A | GACATGACCGTGAAGTTCCATCCGGGCGCAAACC | Reverse PS4 (Major capsid gene *gp033*) |
| **pPF1995** | S14 | Type III-A | GACATGACCGTGAAGTTCCATCCGGGCGCAAACC | Reverse PS5 (Terminase large subunit *gp159*) |
| **pPF1996** | S11 | Type III-A | GTTCTACGACAACAACGAAGCCGGTAACAAAAAC | Reverse PS6 (Tubulin-like gene *gp187*) |
